# Three-step mechanism of promoter escape by RNA polymerase II

**DOI:** 10.1101/2023.12.22.572998

**Authors:** Yumeng Zhan, Frauke Grabbe, Elisa Oberbeckmann, Christian Dienemann, Patrick Cramer

## Abstract

The transition from transcription initiation to elongation is highly regulated in human cells but remains incompletely understood at the structural level. In particular, it is unclear how interactions between RNA polymerase (Pol) II and initiation factors are broken to enable promoter escape. Here we reconstitute Pol II promoter escape *in vitro*, determine high-resolution structures of five transition intermediates, and show that promoter escape occurs in three major steps. First, the growing RNA transcript displaces the B-reader element of the initiation factor TFIIB without evicting TFIIB. Second, rewinding of the upstream edge of the growing DNA bubble evicts TFIIA, TFIIB and TBP and repositions parts of TFIIE and TFIIF. Third, binding of DSIF and NELF evicts TFIIE and TFIIH, establishing the paused elongation complex. This three-step model for promoter escape fills a gap in our understanding of the initiation-elongation transition of Pol II transcription.

## INTRODUCTION

Transcription by RNA polymerase II (Pol II) starts with the formation of the pre-initiation complex (PIC), which consists of Pol II and a set of general transcription factors (GTFs): TFIIA, TFIIB, TFIID, TFIIE, TFIIH and Mediator^1–4^. Upon PIC positioning on the promoter, the translocase activity of TFIIH pumps downstream DNA into Pol II active center cleft and applies torque to induce melting of the DNA duplex^5,6^. As RNA synthesis commences, the PIC transitions to an initially transcribing complex (ITC), which still contains GTFs. When the RNA reaches a certain length, the ITC is converted to an elongation complex (EC) and the transcribing complex escapes from the promoter region^7^.

During promoter escape, the transcription complex undergoes structural and compositional changes^7,8^. As initial transcription proceeds, the upstream end of the initially melted DNA bubble remained in place, whereas the downstream junction of the bubble is further unwound, enlarging the DNA bubble^9,10^. The ITC tends to release the nascent RNA in a process termed abortive transcription^10,11^ and undergoes early promoter-proximal arrest^12–14^. When the transcript reaches a length of 7 nucleotides (nt) or longer, bubble rewinding from the upstream end of the extended bubble leads to the formation of a stable transcription bubble that is consistent with an EC^7,9^. After bubble rewinding, the transcription complex reaches full stability, marked by the end of abortive transcription^10^ and independence of TFIIH translocase activity for efficient elongation^9^.

In addition to the transitions in the DNA and RNA structure during initial transcription, the GTFs are released from Pol II^7,8^. Structural studies of Pol II-TFIIB complexes showed that the TFIIB reader domain projects into the Pol II active center cleft^15^ and would clash with an RNA longer than 7 nt^16^. Structural comparison between Pol II-TFIIB complexes and ECs also showed that the TFIIB linker domain clashes with the upstream DNA after bubble rewinding^15^. These findings led to the early hypothesis that extension of the nascent RNA beyond 6 nt triggers release of TFIIB and promoter escape^15^. However, biochemical studies later showed that TFIIB can stay associated with Pol II until the RNA reaches 12-13 nt^17^. To accommodate TFIIB in the early transcribing complex with a longer RNA, TFIIB needs to rearrange within the transcription complex. However, structural information on such putative conformational changes is lacking.

The fate of TFIIE and TFIIH is also unclear, as different results were reported. For TFIIE, it was suggested that dissociation from Pol II occurs before the RNA transcript reaches 10 nt, whereas TFIIH would leave the early elongation complex after the transcript reaches 30 nt^18^. Another study proposed that TFIIE and TFIIH subunits dissociate in separate steps: TFIIE-α and the TFIIH kinase module dissociate first, whereas TFIIE-β and the core of TFIIH dissociate later^19^. TFIIF has been reported not to be displaced during the initiation-elongation transition but rather to travel with the early EC^18,20^ and to stimulate elongation^21,22^ until being displaced in a casein kinase (CK2) dependent manner^23^. Additionally, the multi-subunit elongation factors Paf1 complex (PAF)^24,25^ and super elongation complex (SEC)^26^ bind to Pol II at sites that overlap with TFIIF binding regions, suggesting their binding is mutually exclusive.

To study the mechanisms of promoter escape, an efficient *in vitro* system for *de novo*, promoter-dependent transcription is required that allows for reconstitution of the initiation- elongation transition in the test tube and for subsequent structural characterization of the transition intermediates. Two recent studies reported such *in vitro* transcription initiation systems for the yeast *Saccharomyces cerevisiae*^27,28^. However, the initiation-elongation transition differs between the yeast and human systems. In particular, yeast Pol II undergoes initial promoter scanning, a process that is absent in human transcription^29–32^, and yeast Pol II does not pause in the promoter-proximal region, a process that occurs in the human system and depends on the negative elongation factor NELF that is absent from yeast^33^.

Here, we describe an experimental setup for highly efficient human *in vitro* promoter- dependent transcription and perform structural investigations of the reaction mixture by cryo- electron microscopy (cryo-EM). We capture five intermediates during human promoter escape and determine their structures. We visualize key steps during promoter escape where the ITC undergoes stepwise structural and compositional changes upon transitioning to an EC. Our results lead to a simple, three-step model for promoter escape by Pol II.

## RESULTS

### Reconstitution of the initiation-elongation transition with human proteins

To reconstitute the initiation-elongation transition of transcription *in vitro*, we assembled the PIC on promoter DNA templates using the human GTFs TBP, TFIIA, -IIB, -IIF, -IIE and –IIH, and *S. scrofa* Pol II, which is 99.9% identical to human Pol II (Figure 1A). As DNA template, we used a variant of the Adenovirus major late promoter (AdMLP) containing a U-less cassette, which allowed us to stall Pol II after synthesis of a 16 nt RNA (Figure 1B top and S1A). Transcription was initiated by the addition of ATP, CTP, GTP, [α-^32^P]-CTP and reactions were incubated at 30°C for 15 minutes.

**Figure 1.**
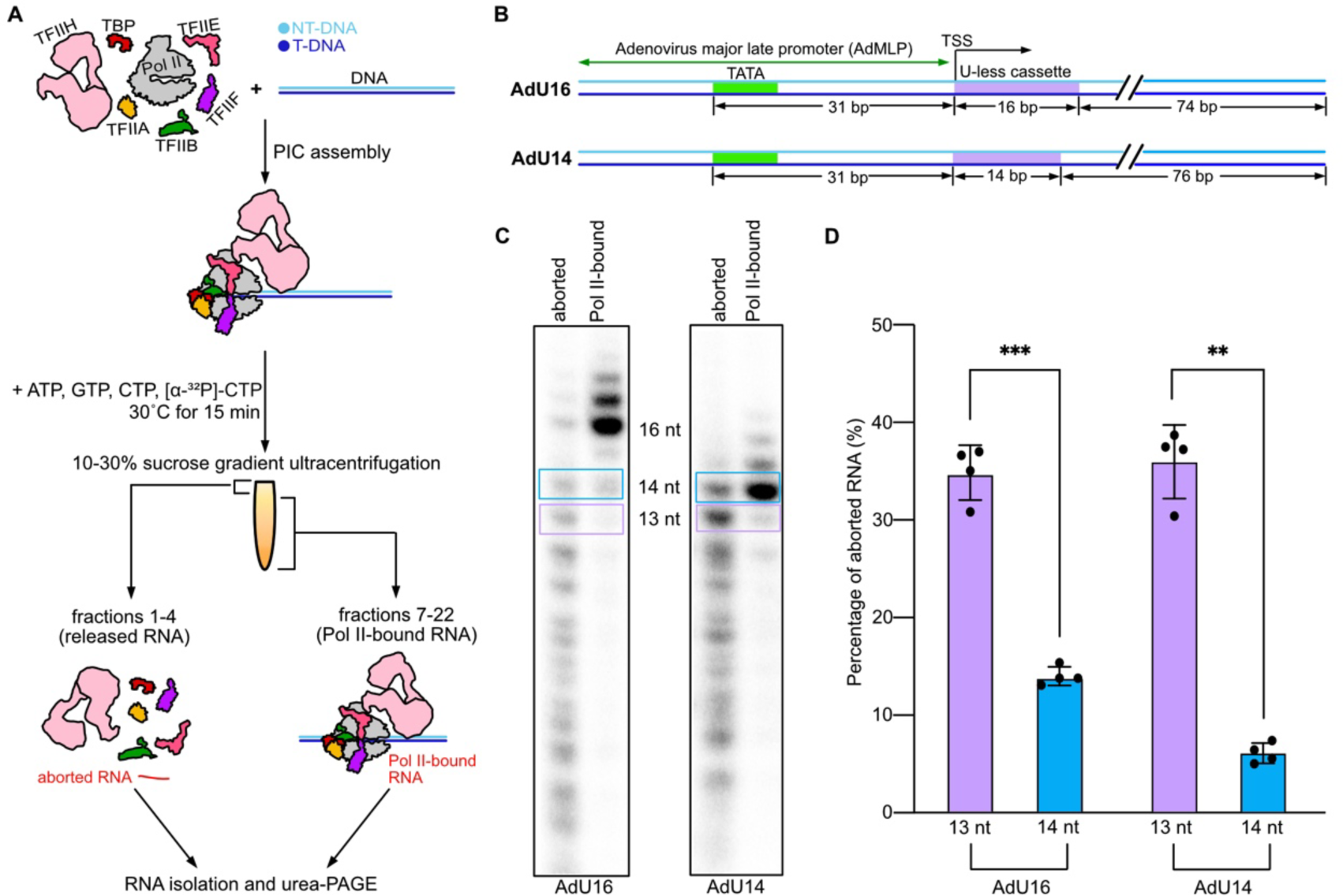
Pol II enters stable RNA chain elongation when nascent RNA is 14 nt long. **A,** Scheme of the experimental setup to measure aborted and Pol II-bound RNA. NT-DNA, non-template DNA; T-DNA, template DNA. **B,** Scheme of the promoter DNA templates used in the experiments. TATA-box and U-less cassettes are highlighted in green and purple, respectively. The lengths of the U-less cassette, the total length of DNA template and the distance between TSS and TATA box are indicated. **C,** Urea-PAGE analysis of aborted and Pol II-bound RNA products from transcription reactions with AdU16 and AdU14 promoter DNAs. Experiments were repeated four times. Rectangle boxes denote bands of interest that were quantified. **D,** Quantification of the fraction of aborted RNAs with lengths of 13 nt and 14 nt from transcription reactions using AdU16 and AdU14 promoter DNA. Mean ± standard deviation (SD) values are indicated. p-values were calculated with Brown-Forsythe ANOVA tests (**p < 0.01 and ***p < 0.001). The color code is the same as panel C. p-values for AdU16 and AdU14 are 0.0007 and 0.0023, respectively.

To monitor abortive transcription, RNA released from Pol II and Pol II-bound RNA were separated by sucrose gradient centrifugation (Figure 1A). The upper factions of the gradient, which are devoid of Pol II, contained excess GTFs and aborted RNA transcripts that were released from Pol II (fractions 1-4). The lower fractions (7-22) contained RNA bound to Pol II (Figure S1B-C). The fractions were pooled and RNAs were resolved by denaturing PAGE (Figure 1C left). We observed released, short abortive RNAs and the full-length 16 nt transcript that was stably associated with Pol II (Figure 1C left and S1D). In summary, we established a highly efficient, promoter-dependent *in vitro* transcription system that enables us to study the transition from initiation to elongation.

When we compared aborted and Pol II-bound RNAs produced by *in vitro* transcription from the AdU16 promoter, we observed a strong reduction of abortive transcription at around 14 nt of RNA (Figure 1C left). We therefore designed the AdU14 promoter that stalls Pol II at 14 nt of RNA (Figure 1B bottom) and resolved aborted and Pol II bound transcripts by denaturing PAGE (Figure 1C). Quantification of RNAs from both the AdU16 and AdU14 promoters showed a strong decrease in the fraction of aborted transcripts when the RNA length grew from 13 nt to 14 nt (Figure 1C-D). This indicates that RNA extension from 13 to 14 nt leads to increased stability of the early transcribing complex. These observations are consistent with previous reports showing that the transcribing complex is unstable and requires TFIIH ATPase activity until the transcript length increases beyond 14 nt^13,14^. Thus, the transition from abortive transcription to stable RNA chain elongation is completed when the RNA reaches 14 nt of length.

### Cryo-EM structures of transition intermediates

We next aimed to structurally characterize early transcription complexes during the transition from abortive transcription to stable elongation by cryo-EM. Guided by our biochemical results, we used the AdU14 promoter that stalls Pol II when the nascent RNA is 14 nt long. We then prepared cryo-EM samples from the transcription reaction (Methods, Figure 2A-B) and collected single-particle cryo-EM data. After extensive data processing, we could refine five structures, three ITC and two EC structures, which were all resolved to ∼3 Å resolution (Figure 2C, S2, S3, S4A-E and Table 1). The high resolution allowed us to assign the nucleic acid sequence in the Pol II active site revealing the exact register of transcription for each complex (Figure S5A). In conclusion, we captured and structurally characterized human transcription intermediates obtained by *de novo* transcription that, in contrast to reconstituted complexes, represent true functional intermediates of the initiation-elongation transition.

**Figure 2.**
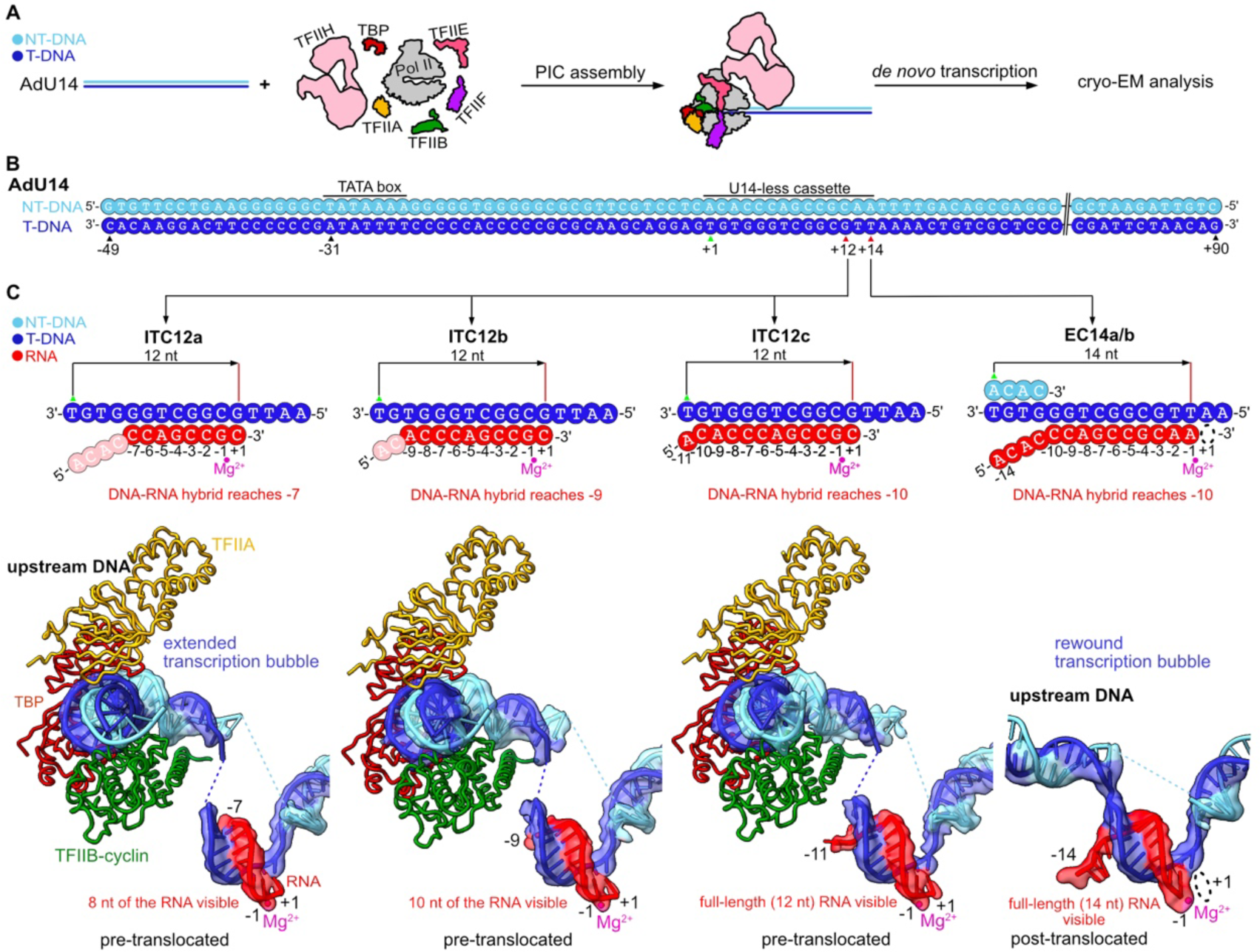
Visualization of early transcription intermediates by cryo-EM. **A,** Scheme of experimental setup for cryo-EM sample preparation. For details, see Methods. **B,** DNA templates used for cryo-EM sample preparation. The TSS (+1) is highlighted by a green triangle. The two stall sites observed in the cryo-EM structures are indicated at +12 and +14 with the register indicated relative to the TSS (+1). **C,** Schematic and cryo-EM density of the five transcription intermediates ITC12a, ITC12b, ITC12c and EC14a/b. Bases of the RNA that were synthesized but structurally disordered are drawn in light red. The cryo-EM densities around the upstream DNA were low-pass filtered to 5 Å for better visualization. The register of the RNA is indicated with respect to the active site. The active site Mg^2+^ is shown as a magenta sphere. For clarity, Pol II and the rest of GTFs are not shown except for TFIIA, TBP and the TFIIB cyclin of ITC12a-c.

**Table 1.** Cryo-EM data collection, refinement and validation statistics.

|  |  |  |  |  |  |
| --- | --- | --- | --- | --- | --- |
| **Data collection and processing** | Dataset1 |  | Dataset2 |  | Dataset3 |
| Magnification | 81,000x |  | 81,000x |  | 81,000x |
| Voltage (kV) | 300 |  | 300 |  | 300 |
| Electron exposure (e<sup>-</sup>/Å<sup>2</sup>) | 40.06 |  | 40.00 |  | 41.47 |
| Defocus range (μm) | 0.7-1.7 |  | 0.7-1.7 |  | 0.7-1.7 |
| Pixel size (Å) | 1.05 |  | 1.05 |  | 1.05 |
| Initial particle images (no.) | 6,386,813 |  | 4,204,757 |  | 2,593,001 |
| **Model name** | ITC12a | ITC12b | ITC12c | EC14a | EC14b |
| PDB code | PDB: YYYY | PDB: YYYY | PDB: YYYY | PDB: YYYY | PDB: XXXX |
| Map code | EMDB: EMD-XXXXX | EMDB: EMD-XXXXX | EMDB: EMD-XXXXX | EMDB: EMD-XXXXX | EMDB: EMD-XXXX |
| Symmetry imposed | C1 | C1 | C1 | C1 | C1 |
| Map resolution (Å) at FSC = 0.143 | 3.1 | 2.9 | 3.4 | 3.4 | 3.3 |
| Map resolution range (Å) | 3.0-11.4 | 2.6-7.9 | 2.8-8.0 | 3.1-10.4 | 3.0-10.6 |
| **Refinement** |  |  |  |  |  |
| Initial models used (PDB code) | 7NW0, 5YID | 7NW0, 5YID | 7NW0, 5YID | 7NW0, 5FLM | 7NW0, 5FLM |
| **Model composition** |  |  |  |  |  |
| Non-hydrogen atoms | 44795 | 44876 | 44728 | 38140 | 36737 |
| Protein residues | 5272 | 5277 | 5249 | 4514 | 4343 |
| Nucleotides | 124 | 127 | 129 | 95 | 95 |
| Ligands | Zn: 10 Mg: 1 | Zn: 10 Mg: 1 | Zn: 10 Mg: 1 | Zn: 9 Mg: 1 | Zn: 9 Mg: 1 |
| **Mean B factors (Å<sup>2</sup>)** |  |  |  |  |  |
| Protein | 116.44 | 116.12 | 125.33 | 151.00 | 144.58 |
| Nucleotides | 171.41 | 157.47 | 198.91 | 201.38 | 199.84 |
| Ligand | 137.62 | 137.69 | 172.74 | 204.85 | 195.06 |
| **R.m.s. deviations** |  |  |  |  |  |
| Bond lengths (Å) | 0.008 | 0.005 | 0.004 | 0.005 | 0.005 |
| Bond angles (°) | 0.839 | 0.671 | 0.698 | 1.176 | 1.168 |
| **Validation** |  |  |  |  |  |
| MolProbity score | 1.36 | 1.45 | 1.67 | 1.88 | 1.87 |
| Clashscore | 3.19 | 3.95 | 5.17 | 9.43 | 8.84 |
| Poor rotamers (%) | 0.00 | 0.00 | 0.00 | 0.00 | 0.00 |
| **Ramachandran plot** |  |  |  |  |  |
| Disallowed (%) | 0.00 | 0.00 | 0.00 | 0.00 | 0.00 |
| Allowed (%) | 3.74 | 3.95 | 5.79 | 5.63 | 5.90 |
| Favored (%) | 96.26 | 96.05 | 94.21 | 94.37 | 94.10 |

When examining the cryo-EM densities for the three ITC structures, we found that they transcribed to register +12 of the AdU14 promoter, and thus, Pol II had stopped transcribing before the designed U-stop at +14. The three ITC structures contain all GTFs and an extended DNA bubble similar to a previously reported ITC structure^34^. The DNA-RNA hybrids in all three ITCs are in the pre-translocated state, however, the number of RNA nucleotides that are visible differs among three ITC structures and thus we called them ITC12a-c (Figure 2C). The two EC structures both contain 14 nt RNAs but showed differences in the composition and conformation of TFIIE (Figure 2C, S4D-F). These structures were called EC14a and EC14b. Both of these structures contain a DNA-RNA hybrid of 10 base pairs (bp) in the post-translocated state and a rewound transcription bubble with an upstream DNA trajectory that is similar to the paused Pol II elongation complex^35^.

### Growing RNA repositions the TFIIB reader but does not displace TFIIB

Although all ITCs have transcribed to +12 of the AdU14 promoter (Figure 2B), we found that in ITC12a and ITC12b only 8 and 10 nt of the RNA are visible (Figure 2C). In ITC12c, however, we could observe density for all 12 nt of the transcribed RNA (Figure 2C). The discrepancy between transcription position and visible RNA length in ITC12a and ITC12b is not caused by the selection of a downstream TSS during *in vitro* transcription because the AdU14 promoter mainly produces RNA of 14 nt and longer (Figure S1E). Therefore, the missing bases at the 5’-end of the RNA in ITC12b and ITC12a were transcribed but are disordered in our cryo-EM reconstructions. As a result, the partially disordered RNA in ITC12a and ITC12b forms shortened hybrids with the DNA template that only reach register -7 and -9, respectively (Figure 2C and 3). In contrast, the RNA in ITC12c forms a hybrid until register -10, which represents a fully formed DNA-RNA hybrid before strand separation at register -11 (Figure 3). These observations suggests that during initial transcription, the DNA-RNA hybrid within the ITC can retain some mobility.

**Figure 3.**
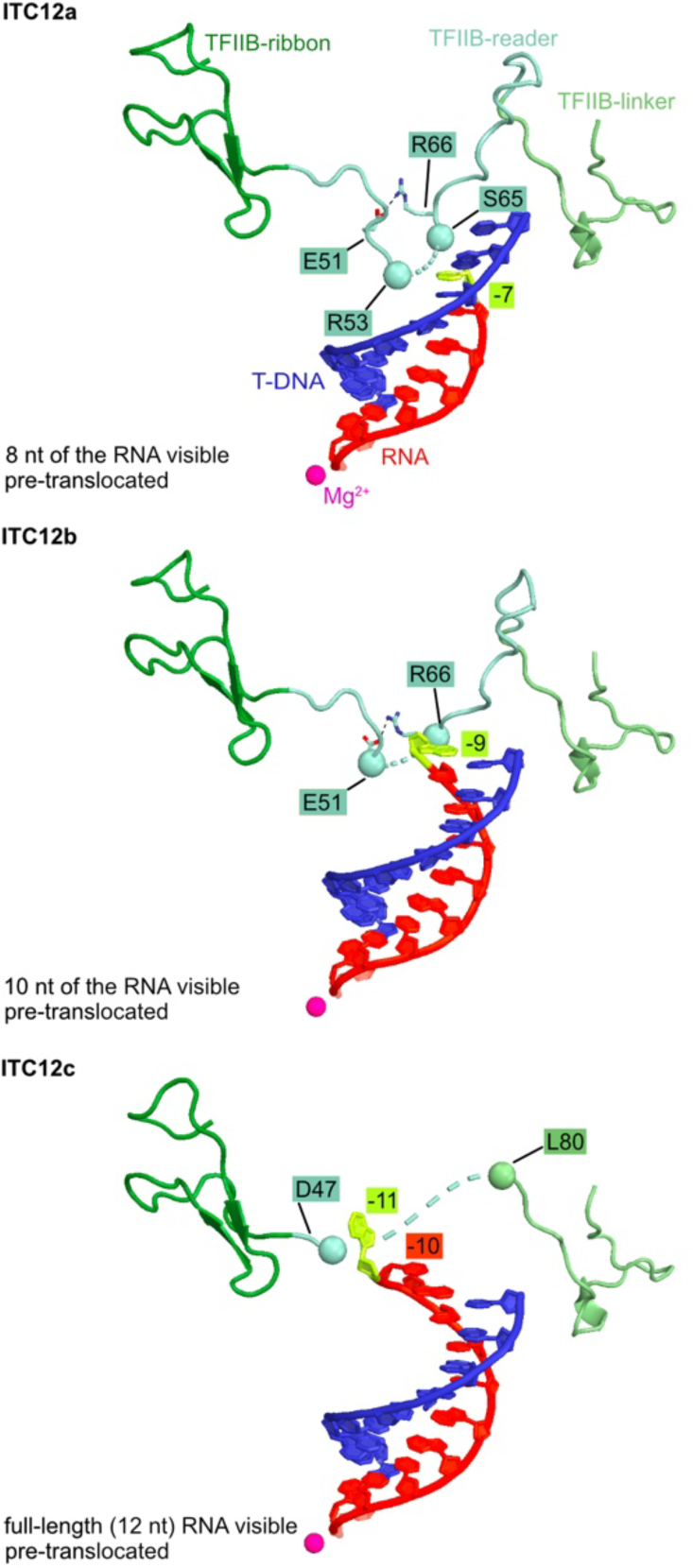
The TFIIB reader competes with the nascent RNA during initial transcription. TFIIB and the DNA-RNA hybrid of ITC12a-c are shown as cartoon. The salt bridge formed by E51 and R66 is indicated by black dashes. The boundaries of the structured parts of TFIIB are shown as spheres. The first visible base at 5’-end of the RNA is highlighted in lemon and its position relative to the active site (-1) is indicated. The active site Mg^2+^ is shown as a magenta sphere. The template DNA at Pol II active site is shown in blue. For clarity, the rest of the DNA, TFIIB- cyclin, Pol II and the other GTFs are omitted.

Since the 5’-end of the initially transcribed RNA interacts with the reader domain of TFIIB^16^, we compared the conformations of the B-reader among the three ITC12 structures (Figure 3 and S5B-D). In ITC12a, residues 54-64 of the B-reader are disordered, indicating that the DNA-RNA hybrid reaching to register –7 displaced this part of the B-reader. When the hybrid reaches register –9 in ITC12b, B-reader residues 52-65 are additionally displaced. Upon formation of a full DNA-RNA hybrid in ITC12c, we could not observe density for residues 48-79 of the TFIIB reader domain. Based on these observations we conclude that the RNA strand of the DNA-RNA hybrid and the B-reader directly compete for the same space in the active center cleft of Pol II and can displace each other as the RNA is growing during early transcription. We suggest that if the B-reader outcompetes most of the RNA from the DNA-RNA hybrid, the initially transcribed RNA may dissociate as a short abortive transcript. However, when the RNA stably forms a full DNA-RNA hybrid, the B-reader is displaced and initial transcription proceeds.

Consistent with this, it was shown that early transcription complexes are prone to stall when the RNA reaches a length of 7-9 nt^9,36^ and that this stalling can be eliminated by the TFIIB point mutation R66L, which destabilizes a conserved salt bridge within the B-reader^9^. Modelling based on our structures suggests that formation of a full 10 nt DNA-RNA hybrid requires disruption of the stabilizing salt bridge formed between E51 and R66 (Figure 3). This provides a possible mechanism for how the B-reader and RNA compete – salt bridge formation stabilizes the B-reader whereas the growing RNA destabilizes the salt bridge and thereby also the B-reader.

### Bubble rewinding and RNA extension evict TBP, TFIIA and TFIIB

As RNA synthesis proceeds, the extended, early DNA bubble rewinds from upstream, and this rewinding was suggested to coincide with the transition from abortive to stable RNA synthesis^9^. Consistent with this model, we observe a strong difference in upstream DNA when we compare our ITC with our EC structures. Whereas the ITC structures that transcribed 12 nt of RNA all contain extended upstream bubbles, our EC structures show that upstream DNA is rewound and positioned where it is observed in a reconstituted paused elongation complex^35^.

To investigate the structural changes within the transcription complex upon bubble rewinding, we compared the structures of ITC12c and EC14a. In EC14a, we do not observe density for the upstream promoter complex comprising TBP, TFIIA and TFIIB (Figure 4 and S4D- E). Superposition of the ITC with the EC structures shows that the rewound upstream DNA duplex clashes with TFIIB cyclin and linker domains (Figure S4G). In the ITC, TBP and TFIIA are mainly binding to the DNA and the TFIIB cyclin domain (Figure S4G). Therefore, bubble rewinding and displacement of TFIIB from Pol II also evicts TBP and TFIIA. These observations suggest that upstream DNA rewinding leads to eviction of the upstream complex from Pol II, whereas the B-ribbon is displaced by the 14 nt long transcript, which starts clashing with the B-ribbon at a length of 12 nt (Figure S4H). This is consistent with a previous study showing that TFIIB is only destabilized when the RNA is 12-13 nt long^17^. In summary, RNA extension and DNA bubble rewinding lead to eviction of TBP, TFIIA and TFIIB, and convert the ITC to an early EC.

**Figure 4.**
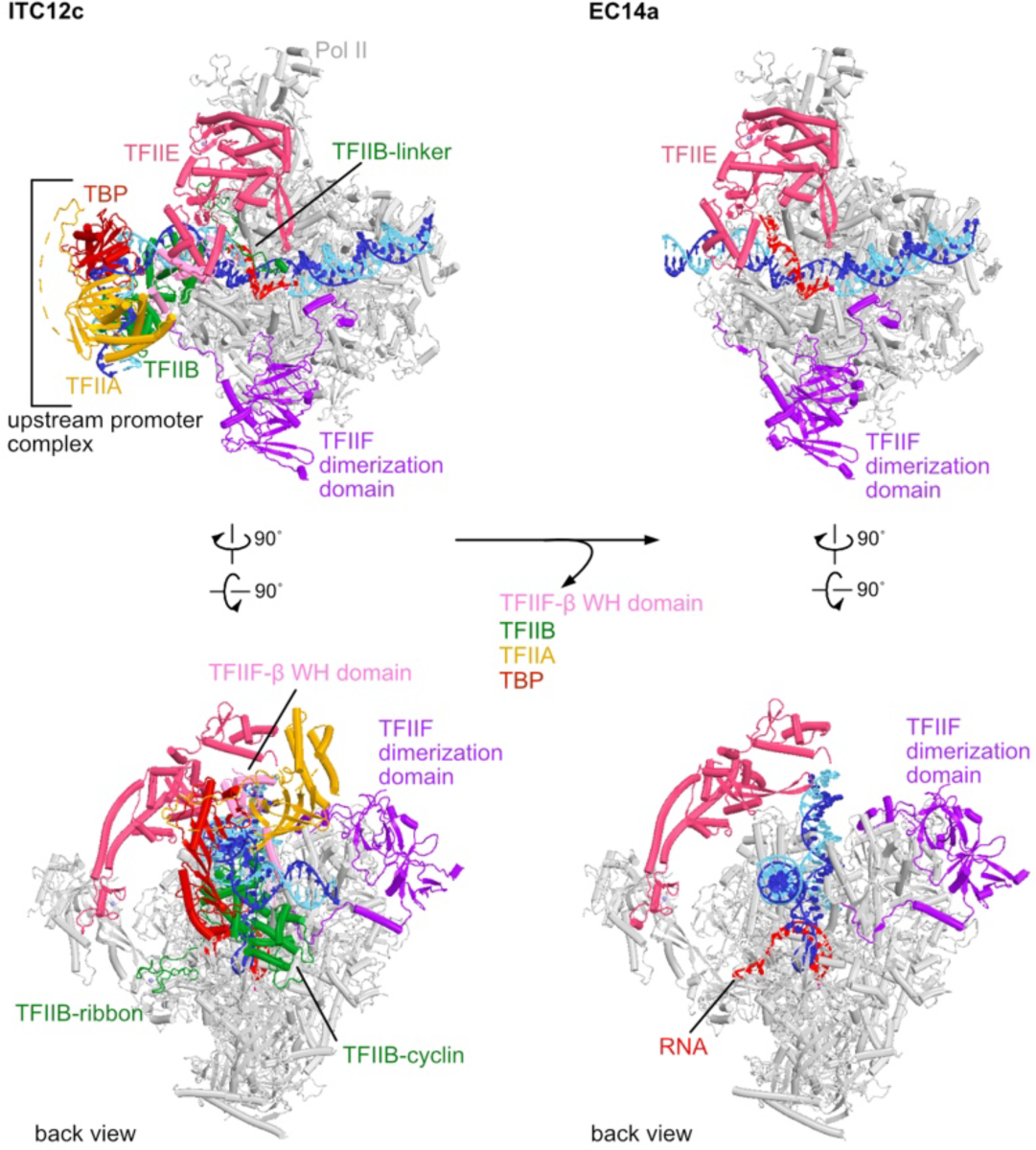
Bubble rewinding and RNA extension evict TBP, TFIIA and TFIIB. Structures of ITC12c (left) and EC14a (right) are shown as cartoon for comparison.

### TFIIF, TFIIE and TFIIH remain bound to the early elongation complex

It was proposed that bubble rewinding during the conversion from the ITC to the early EC also triggers release of TFIIE and TFIIH^7,33^. In contrast to that suggestion, we observe density for TFIIF, TFIIE and TFIIH in our cryo-EM reconstructions of both ECs (Figure 5A-B). TFIIF is bound to Pol II in both ECs but we could not observe density for the TFIIF-β winged-helix (WH) domain (Figure 4). The TFIIF-β WH domain serves as a binding site for TFIIE-β in the PIC and ITC, and dissociation of the WH could influence the conformation and occupancy of TFIIE within the early EC.

**Figure 5.**
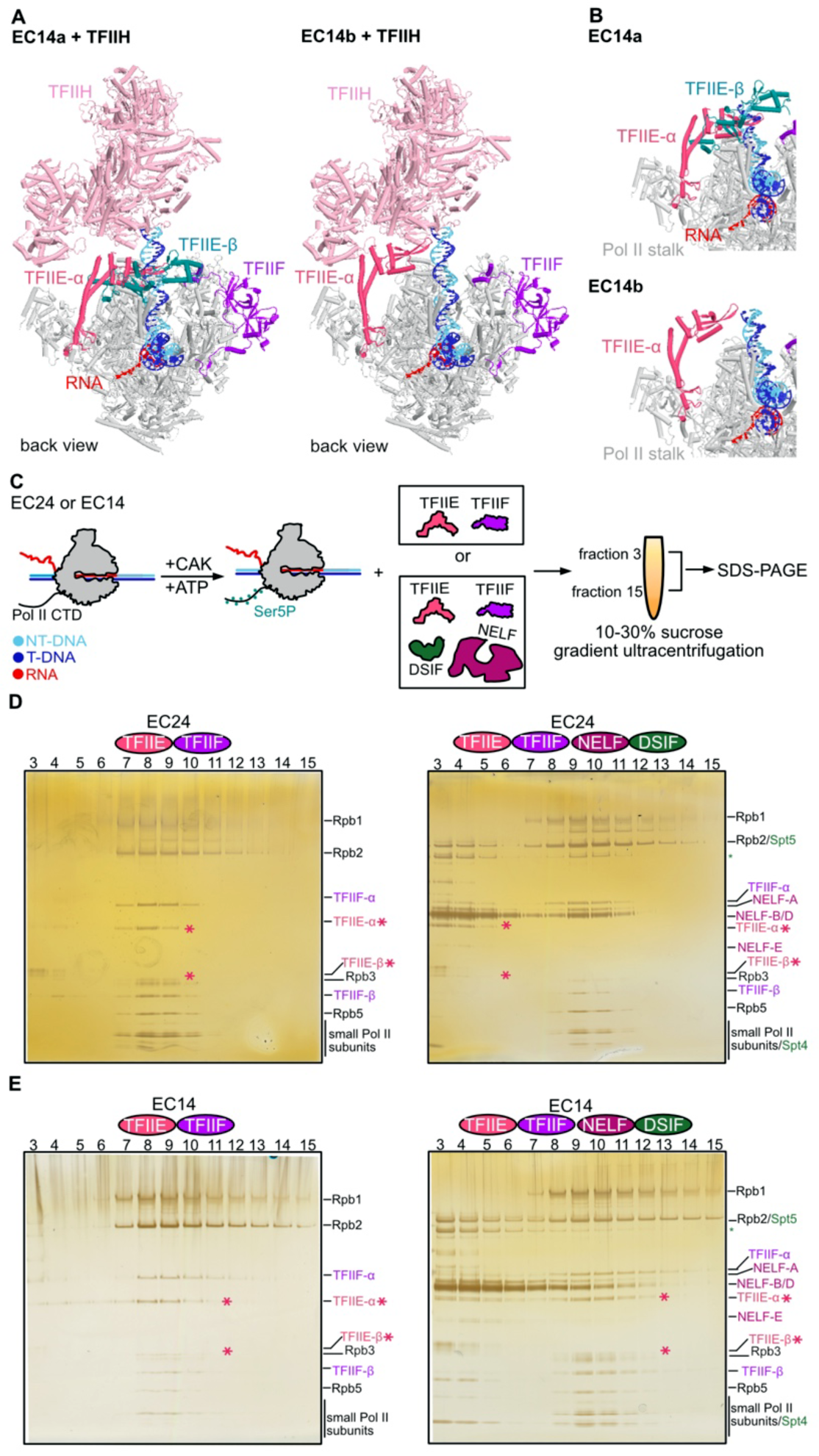
TFIIE, TFIIF and TFIIH remain bound to the early elongation complex. **A,** Structures of EC14a and EC14b containing TFIIH, TFIIE and TFIIF. The structure of TFIIH (PDB: 7nvx^38^) was rigid-body docked into the global refined maps of EC14a+TFIIH and EC14b+TFIIH (Methods). **B,** Two states of TFIIE-α and -β subunits in the early elongation complex. Close-up views of TFIIE and the Pol II stalk in EC14a (top) and EC14b (bottom) are shown. **C,** Scheme of the experimental setup to monitor the displacement of TFIIE by DSIF and NELF. For details, see Methods. **D,** Silver-stained SDS-PAGE analysis of peak fractions of EC24 after analytical gradient ultracentrifugation. Protein bands corresponding to TFIIE are highlighted with pink asterisks. Green asterisk marks a degradation band of the DSIF subunit Spt5. **E,** Silver-stained SDS-PAGE analysis of peak fractions of EC14 after analytical gradient ultracentrifugation. Protein bands corresponding to TFIIE are highlighted with pink asterisks. Green asterisk marks a degradation band of the DSIF subunit Spt5.

Indeed, when we classified the particle populations of the ECs after masking the region around TFIIE, we observed two distinct conformations of TFIIE (Figure 5B, S2, S3 and S4D-F). In EC14a, density for both subunits of TFIIE is visible. However, in EC14b, clear density is only present for the TFIIE-α subunit, which additionally has rotated around the Pol II stalk (Figure S4F).

TFIIE-α serves the main interaction partner for TFIIH within the ITC^34,37–39^ and TFIIE is crucial for binding of TFIIH to the PIC^39^. In line with this, we find that TFIIH only binds the early EC in the presence of TFIIE (Figure S6A-B) and we find density for TFIIH in the cryo-EM reconstructions of both EC14a and EC14b (Figure 5A). In summary, TFIIF, TFIIE and TFIIH remain bound to the early elongation complex, demonstrating that neither RNA extension nor upstream DNA rewinding can trigger their release from the early EC.

### DSIF and NELF displace TFIIE and TFIIH

From these results, it remained unclear how TFIIF, TFIIE and TFIIH are removed from the early EC. We therefore asked whether the general elongation factor DSIF, the capping enzyme RNGTT or the pausing factor NELF, which bind when RNA reaches lengths of 24-50 nt^33,40^ may be responsible for eviction of the remaining GTFs.

To this end, we first assembled an EC24 complex with TFIIF, TFIIE and a 24 nt RNA (Methods, Figure 5C). TFIIH was not included in the experiments as its binding to the early EC is strictly dependent on TFIIE and displacement of TFIIE will therefore inevitably result in dissociation of TFIIH as well. Although TFIIF is likely displaced by elongation factors^26,41–43^, we still included TFIIF in the experiments as it is present in the early EC (Figure 5A) and may interact with TFIIE and stabilize it. We then added DSIF together with the capping enzyme RNGTT, but could not observe eviction of TFIIF or TFIIE (Figure S6C-D). However, when adding DSIF in combination with NELF, TFIIE was completely displaced from the EC24, whereas TFIIF remained associated (Figure 5D and S6E left). DSIF and NELF however failed to displace TFIIE from an EC14 with 14 nt of RNA (Figure 5E and S6E right), in line with previous studies showing that the binding of DSIF and NELF requires an RNA of at least 22 nt length^44^. We conclude that binding of DSIF and NELF evicts TFIIE from the early EC, and that this relies on RNA extension beyond 22 nt. Additionally, we suggest that TFIIH is leaving the early EC when TFIIE dissociates, because TFIIE is strictly required for TFIIH binding. In summary, we conclude that TFIIE and TFIIH can remain associated with the early EC until RNA growth recruits DSIF and NELF.

## DISCUSSION

Here, we establish an *in vitro* promoter-dependent human transcription system that enables us to carry out structural studies of early Pol II complex intermediates and to investigate the initiation-elongation transition of transcription. Our results converge with published data on a three-step model of promoter escape by Pol II (Figure 6, Movie 1). First, when Pol II synthesizes an RNA longer than 6 nt, the transcript begins to compete with the TFIIB reader domain for the same binding site in the Pol II active center, which may cause Pol II stalling and abortive transcription. As the RNA extends, it can however displace the B-reader without evicting TFIIB. Second, when the RNA reaches a length of 14 nt, the upstream DNA bubble collapses and rewinding of upstream DNA leads to eviction of the upstream promoter complex comprising TBP, TFIIA, TFIIB and displacement of the TFIIF-β WH domain from the Pol II surface. Third, when the RNA extends to at least 22 nt and the elongation factors DSIF and NELF are recruited, binding of these factors to the Pol II surface displaces TFIIE and TFIIH. Finally, it is known that other elongation factors bind the Pol II lobe region and can displace TFIIF^26,41–43^. Displacement of all GTFs from transcribing Pol II marks the end of promoter escape and the transition to an EC.

**Figure 6.**
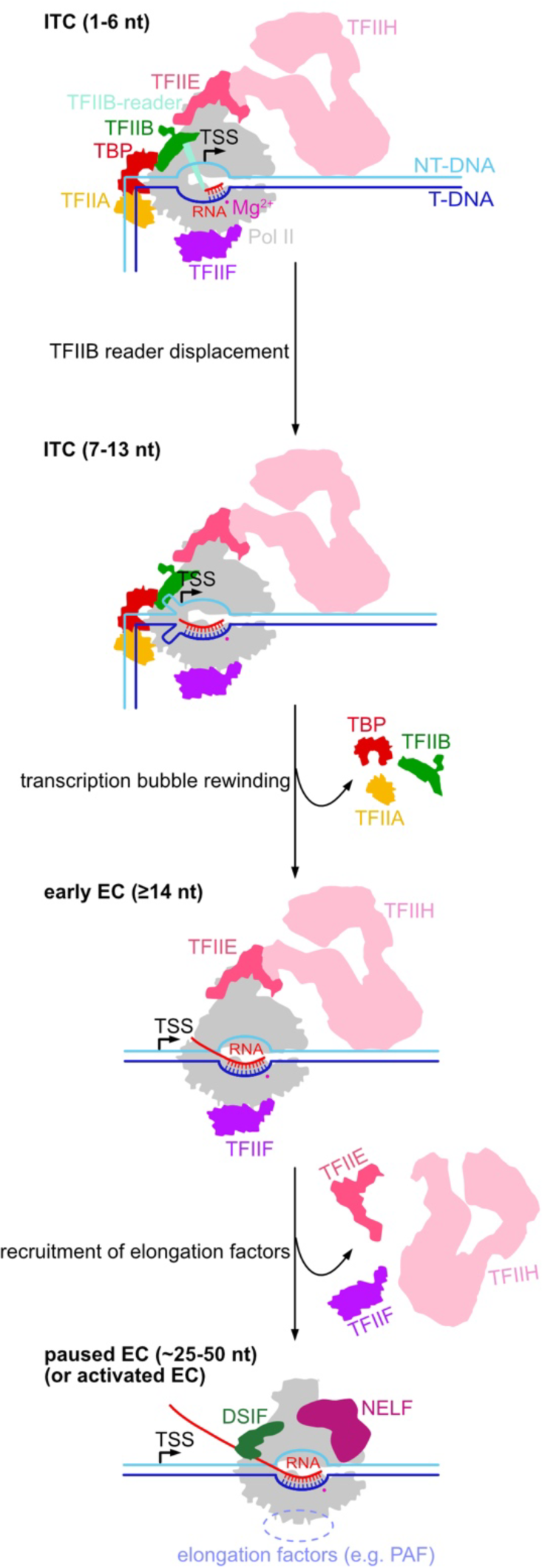
Three-step model for promoter escape by human Pol II. Promoter escape by human Pol II occurs in three steps. First, the RNA transcript competes with and eventually displaces the TFIIB reader from the Pol II active center when the RNA is around 7-13 nt long. Second, rewinding of the upstream transcription bubble leads to eviction of the upstream promoter complex (TBP, TFIIA and TFIIB) from Pol II. Finally, when the RNA is around 25-50 nt^33^, DSIF and NELF are recruited to the early EC and displace TFIIE and TFIIH. Association of elongation factors like PAF likely displace TFIIF^26,41–43^.

Our three-step model generally agrees with previous models, but also differs in one important way. Namely, it was previously proposed that removal of all GTFs from the ITC is triggered by rewinding of the initial extended transcription bubble^7,33^. However, the comparison of our ITC and EC structures, which were formed by *de novo* transcription, strongly suggests that bubble rewinding only leads to dissociation of the upstream promoter complex comprising TBP, TFIIA and TFIIB, whereas the three other GTFs remain bound to Pol II at this stage. Instead, we found that DSIF and NELF can dissociate TFIIE and TFIIH. We speculate that TFIIE and TFIIH may remain associated with the early EC to enable complete Ser5 phosphorylation on the Pol II C-terminal repeat domain (CTD) by the TFIIH kinase CDK7, possibly until high levels of CTD phosphorylation release Mediator^37,45^, which allows recruitment of DSIF and NELF.

Our observation that TFIIF is part of the early EC is consistent with reports that TFIIF can remain bound to the early EC after the initiation-elongation transition^18,20^ where it likely stimulates early transcription elongation^21,22^. However, TFIIF is largely absent from gene bodies^46–48^ and therefore must be displaced soon after the initiation-elongation transition. It was suggested that the activated EC (EC*) is incompatible with TFIIF binding because a region of the PAF complex subunit Leo1 occupies a similar site on the Pol II protrusion as a part of TFIIF^24,25^ (Figure S7A). Together with our observation that TFIIF remains bound to the early EC and that it is not displaced in the context of a paused Pol II elongation complex (PEC) containing DSIF and NELF (Figure 5C-D and Figure S7B), this suggests that TFIIF might get evicted upon the conversion of the PEC to the EC*. Alternatively, Gdown1^43^, components of the SEC^26^ (Figure S7C) or CK2^23^ could release TFIIF. Further studies about the interplay of these factors and the PEC will help to elucidate TFIIF displacement further.

Comparison with the bacterial RNA polymerase (RNAP) system shows that promoter escape involves conceptually similar steps, although the involved proteins strongly differ. First, a short RNA is synthesized that starts clashing with the σ3.2 linker in the initiation factor σ upon extension beyond 5-6 nt^49^. The steric presence of the σ3.2 linker in the path of the newly synthesized RNA was shown to be the cause for abortive transcription by RNAP^49,50^. The magnitude of abortive transcription is highly dependent of the sequence of the initially transcribed region^51^. Rewinding of the initial transcription bubble triggers partial dissociation of the σ factor^52^. Displacement of the σ factor can be achieved by binding of the elongation regulatory factor NusA^53^, which stabilizes promoter-proximally paused RNAP^54^.

With respect to the archaeal transcription system, the RNA polymerase resembles Pol II and archaea contain the initiation factors TBP, TFB and TFE that are related to human TBP, TFIIB and TFIIE, respectively^55^. Although promoter escape has not been studied in detail in this system, it has been shown that the TFB reader stimulates abortive transcription^56^ and is displaced when the RNA extends to 10 nt^57^. This precedes the complete release of TFB which only occurs when the RNA is longer than 15 nt^57^. Additionally, it is known that binding of the DSIF-related archaeal elongation factor Spt4-Spt5 occurs at the polymerase clamp^58^ and can displace the archaeal TFIIE counterpart TFE^59^. Thus, although details differ, present evidence strongly indicates that the three key steps of promoter escape are conserved from bacteria to eukaryotes. We suggest that the conserved mechanism of promoter escape is likely a consequence of the nature of nucleic acid conformation, interaction and energetics, whereas different proteins evolved around DNA and RNA to catalyze transcription initiation and the transition to elongation.

## Limitations of the study

Due to the highly dynamic nature of promoter escape and the flexibility of transcription complexes, extensive classification of the cryo-EM images was performed to sort out the conformational and compositional heterogeneity of the particles. In turn, certain states might be filtered out during data processing due to their low occurrence. Additionally, we were not able to resolve TFIIH at high resolution, likely due to the presence of ATP during sample preparation. This limits our understanding on the mechanism of TFIIH during promoter escape. In the future, it may be possible to include additional protein factors, to optimize sample preparation and to stabilize TFIIH within certain transcription complex intermediates.

## ACKNOWLEDGEMENTS

We thank current and past members of the Cramer laboratory, in particular, U. Steuerwald for maintenance of the electron microscopy facility, C. Oberthür for purifying DSIF and NELF, G. Garg for providing RNGTT, and G. Kokic, J. Abril-Garrido, J. Walshe and S. Schilbach for discussions; O. Dybkov for providing the glass plates for RNA sequencing gel and advice on radioactivity experiments.

## AUTHOR CONTRIBUTIONS

Y.Z. designed and carried out all experiments and data analysis, unless stated otherwise. Y.Z. and C.D. collected cryo-EM data. F.G. purified general transcription factors. E.O., C.D. and P.C. supervised research. Y.Z., E.O. and C.D interpreted the data. Y.Z., C.D. and P.C. wrote the manuscript with input from all authors.

## DECLARATION OF INTERESTS

The authors declare no competing financial interest.

## STAR METHODS

### Resource Availability Lead contact

Correspondence and request for materials should be addressed to C.D.

### Material Availability

Materials are available from Christian Dienemann upon request under a material transfer agreement with the Max Planck Society.

### Data and Code Availability

The cryo-EM reconstructions and final models were deposited with the EM Data Base (accession code: EMD-XXXXX) and with the Protein Data Bank (accession code: PDB-YYYY).

## METHODS DETAILS

### Cryo-EM sample preparation with AdU14 promoter DNA

AdML promoter scaffolds containing U-less cassettes of different lengths were inserted into pUC119 vectors as previously described^38^. DNA templates were amplified by large-scale PCR, purified by ion-exchange chromatography with a ResourceQ column (5 mL, Cytiva) followed by phenol-chloroform extraction. The sequences of the DNA templates used for cryo-EM sample preparation are listed below. TATA-box and TSS are highlighted in bold. U-less cassettes are underlined. AdU14 template (139 bp): 5’-GTG TTC CTG AAG GGG GGC **TAT AAA A**GG GGG TGG GGG CGC GTT CGT CCT C**<u>A</u>**<u>C ACC CAG CCG CAA</u> TTT TGA CAG CGA GGG CCA GCA GAA GGG GAG AGA ACT TTT TCT CAA AAG CGG GCA TGA CTT CTG CGC TAA GAT TGTC- 3’.

*S. scrofa* Pol II and human initiation factors (TBP, TFIIA, TFIIB, TFIIE, TFIIF and TFIIH) were purified as previously described^35,38,60–62^. Sample preparation was performed based on published protocol^38^ with modifications. Before PIC formation, three complexes, DNA-TFIIA-TFIIB-TBP, Pol II-TFIIF and TFIIH-TFIIE were incubated as three separate samples at room temperature for 5-10 min. Then, the pre-assembled samples were mixed and further incubated at 30°C for 1 h. Transcription was initiated by the addition of 0.5 mM NTP mix consisting of ATP, CTP and GTP. Transcription was allowed to proceed for 2∼10 min at 30°C in reaction buffer containing 20 mM K-HEPES pH 7.5, 110 mM KCl, 3% glycerol, 16 mM MgCl2, 0.5 mM DTT. Afterwards, 6/7^th^ of the resulting transcription reaction was loaded onto a 15%-40% (w/v) sucrose gradient while mildly cross-linked by GraFix^63^ at 32,000 rpm (SW60 rotor) for 16 h at 4 °C. After centrifugation, the gradient was fractionated manually from the top and quenched with 10 mM lysine and 40 mM aspartate for 10 min on ice. In parallel to GraFix, the 1/7^th^ of the same transcription reaction was loaded onto a 15%-40% (w/v) sucrose gradient but without cross-linker to monitor the distribution of the transcription complexes. After ultracentrifugation, fractions from non-crosslinked sample were isopropanol precipitated and analyzed by urea-PAGE. Corresponding fractions that contained the transcribed complexes from the cross-linked sample were then pooled and dialyzed against the Buffer D (20 mM K-HEPES pH 7.5, 90 mM KCl, 1% glycerol (v/v), 1mM DTT) at 4°C for 7-8 h to remove sucrose and glycerol. Subsequently, a thin piece of homemade continuous carbon film (∼2.7 nm) was floated onto the dialyzed sample and the particles were allowed to adsorb onto the film at 4°C for 15 min. The film was then picked up by a holey-carbon grid (Quantifoil R3.5/1, copper, mesh 200) and 3.5 μL of Buffer D was immediately added onto the carbon film prior to vitrification with a Vitrobot Mark IV (FEI) operated at 4 °C and 100% humidity. The grids were blotted for 1-1.5 s with blot force of 5 before plunge-freezing in liquid ethane.

### Visualization of abortive transcription

Abortive transcription was visualized by gradient ultracentrifugation. For that, promoter dependent *in vitro* transcription reactions were performed with AdU14 and AdU16 promoter DNA. The sequences of AdU14 templates were same as cryo-EM sample preparation. The sequence of AdU16 template is listed below. TATA box and TSS are highlighted as bold. U16-less cassette is underlined. AdU16 template (139 bp): 5’-GTG TTC CTG AAG GGG GGC **TAT AAA A**GG GGG TGG GGG CGC GTT CGT CCT C**<u>A</u>**<u>C ACC CAG CCG CAA CG</u>T TTT CAG CGA GGG CCA GCA GAA GGG GAG AGA ACT TTT TCT CAA AAG CGG GCA TGA CTT CTG CGC TAA GAT TGTC- 3’.

PIC assembly was carried out in the same way as sample preparation. For each 20 µL reaction, 14 pmol Pol II, 71 pmol TFIIF, 15 pmol DNA template, 93 pmol TFIIA, 47 pmol TBP, 47 pmol TFIIB, 21 pmol TFIIE and 21 pmol TFIIH were used. Transcription was initiated with 0.5 mM ATP, 0.5 mM CTP, 0.5 mM GTP and 2 µCi [α^32^P]-CTP and allowed to take place for 15 min at 30°C in reaction buffer containing 20 mM K-HEPES pH 7.5, 110 mM KCl, 3% glycerol, 16 mM MgCl2, 0.5 mM DTT. Transcription reactions were then loaded onto a 10%-30% (w/v) sucrose gradient in buffer containing 20 mM K-HEPES pH 7.5, 110 mM KCl, 3% glycerol (v/v), 1 mM DTT, 0.5 mM MgCl2. Ultracentrifugation was performed at 55,000 rpm for 3 h at 4°C with a swinging bucket rotor S55S (Thermo Scientific). The gradients were fractioned manually from top. Fractions 1-4 and 7-22 were pooled separately and digested with proteinase K in the presence of GlycoBlue (Thermo Fisher) for 30 min at 37°C followed by isopropanol precipitation. The precipitated RNA pellets were resuspended in 2x RNA loading dye (NEB) and loaded onto an RNA sequencing gel (7 M urea, 1x TBE, 20% acrylamide:bis-acrylamide 19:1). To achieve similar exposure of the phosphorus screen, 2/15^th^ of the RNA suspension from fraction 7-22 were loaded whereas the total RNA suspension from fraction 1-4 was loaded. For quantification see below.

### Competition assay of TFIIE and DSIF, NELF and RNGTT on early ECs

Analytical gradient ultracentrifugation was used to monitor the competition between TFIIE and elongation factors. DSIF, RNGTT and NELF were purified as previously reported^35,40,64^. To assemble the Pol II ECs, the template DNA strand was mixed with RNA in equimolar ratio and annealed by first incubating at 75°C for 1 min then slowly cooling down to 4°C at the speed of 1°C per min. Next, the DNA-RNA scaffold was added to Pol II in equimolar ratio and incubated for 10 min at 30°C. Finally, the non-template DNA strand was added in 2x molar excess and incubated for another 10 min at 30°C. DNA and RNA oligonucleotides were purchased from Integrated DNA technologies. Sequences used for EC14 assembly were: template: 5′-TTC TGC TGG CCC TCG CTG TCA AAA TTG CGG CTG GGT GTG AGG ACG AAC GCG CCC CCA CCC CCT TTT ATA GCC CCC CTT CAG GAA CAC-3′; non-template: 5′- GTG TTC CTG AAG GGG GGC TAT AAA AGG GGG TGG GGG CGC GTT CGT CCT CAC AGG GTC GGC GTT TTT TGA CAG CGA GGGCCA GCA GAA-3′; RNA: 5’-rUrUrU rCrCrC rArGrC rCrGrC rArA-3’. Sequences used for EC24 assembly were: template: 5′-GAC AAT CTT AGC GCA GAA GTC ATG CCC GCT TTT GAG AAA AAG TTC TCT CCC CTT CTG CTG GCC-3′; non-template: 5′- GGC CAG CAG AAC CCC TCT CTT GTT TTT CTC AAA AGC GGG CAT GAC TTC TGC GCT AAG ATT GTC -3′; RNA: 5’-rUrCrC rCrGrG rUrCrG rUrCrU rUrGrG rGrGrA rGrArG rArArC-3’. The preformed EC was then incubated with CAK (1 µM) and ATP (1 mM) for 30 min at 30 °C to phosphorylate the Pol II CTD in final buffer containing 20 mM K-HEPES pH 7.5, 100 mM KCl, 4% glycerol, 4 mM MgCl2 and 1 mM DTT. The phosphorylation reaction was stopped with 8 mM EDTA before adding initiation factors (TFIIE and TFIIF) in 2x molar excess and incubated at 30°C for 15 min. Afterwards, elongation factors (DSIF and NELF or DSIF and RNGTT) were added in 2x molar excess and incubated at 30°C for 15 min. Reactions were loaded onto a 10%-30% (w/v) sucrose gradient in buffer containing 110 mM KCl, 20 mM HEPES (pH 7.5), 3% glycerol (v/v), 1 mM DTT, 0.5 mM MgCl2. Ultracentrifugation was performed at 32,000 rpm for 16 h at 4°C using an SW60 rotor. Gradient fractions were analyzed by SDS-PAGE.

### TFIIE-dependent TFIIH binding to the early EC

To monitor TFIIE-dependent TFIIH association with Pol II by gradient ultracentrifugation, the two GTFs were added to a preformed EC14 in 2x molar excess and incubated at 30°C for 30 min in final buffer containing 20 mM K-HEPES pH 7.5, 100 mM KCl, 4% glycerol, 4 mM MgCl2 and 1 mM DTT. EC14 was formed as described above but not phosphorylated by CAK. Reactions with and without TFIIE were loaded onto a 10%-30% (w/v) sucrose gradient in buffer containing 110 mM KCl, 20 mM K-HEPES pH 7.5, 3% glycerol (v/v), 1 mM DTT, 0.5 mM MgCl2. Ultracentrifugation was performed at 32,000 rpm for 16 h at 4°C with SW60 rotor. Fractions were analyzed by SDS-PAGE.

### *In vitro* transcription assay

Promoter-dependent *in vitro* transcription assays were performed as previously described^38^, albeit with modifications. In brief, PIC assembly was carried out in the same way as cryo-EM sample preparation. For each 10 µL reaction, 5 pmol Pol II, 25 pmol TFIIF, 5.3 pmol DNA template, 34 pmol TFIIA, 17 pmol TBP, 17 pmol TFIIB, 8 pmol TFIIE and 8 pmol TFIIH were used. Transcription was initiated with 0.5 mM ATP, 0.5 mM CTP, 0.5 mM GTP and 2 µCi [α^32^P]-CTP and allowed to take place for 15 min at 30°C in reaction buffer containing 20 mM K-HEPES pH 7.5, 110 mM KCl, 3% glycerol, 16 mM MgCl2, 0.5 mM DTT. Transcription was stopped by the addition of 2 µg proteinase K (New England Biolabs), 50 mM EDTA, 300 mM sodium acetate and incubated at 37°C for 30 min. RNA was isopropanol-precipitated in the presence of GlycoBlue (Thermo Fisher) overnight at -20°C. The precipitated RNA pellets were resuspended in 2x RNA loading dye (NEB) and loaded onto an RNA sequencing gel (7 M urea, 1x TBE, 20% acrylamide:bis-acrylamide 19:1).

### Cryo-EM data collection and processing

Cryo-EM data were collected on a FEI Titan Krios transmission electron microscope with a K3 summit direct electron detector (Gatan) and a GIF quantum energy filter (Gatan), operated at 300 keV and with a slit width of 20 eV. Data collection was performed automatically with SerialEM^65^ at a nominal magnification of x81,000 (1.05 Å per pixel) with a total dose of around 40 e/Å^2^ fractionated over 40 frames. The defocus range applied was 0.7 µm to 1.7 µm.

On-the-fly motion correction, contrast-transfer function estimation and particle picking were performed with Warp^66^. Initial cleaning of the datasets was carried out by consecutive rounds of 2D and 3D classification in cryoSPARC^67^ to remove ice contamination, falsely picked and aggregated particles. Further processing steps were performed in Relion3.1^68,69^, as described in detail in Figure S2. Focused 3D classification with a mask around core Pol II was performed to separate complexes with open and closed DNA. Particles that contained high resolution features for the DNA-RNA hybrid in the Pol II active site were CTF-refined and polished. These particles were further signal subtracted and classified with a mask around the upstream DNA to separate complexes before and after bubble rewinding. Particles with features of extended bubble were further classified with a mask around core Pol II to separate ITCs with different DNA-RNA hybrids. Particles with features of a rewound bubble were further signal subtracted and classified with a mask around TFIIE to separate different TFIIE conformations. Focused refinements were performed to improve the resolution of Pol II in each class.

Particles with high-resolution features for DNA-RNA hybrid were also signal subtracted and classified with a mask around TFIIH. Particles with concrete features of TFIIH were combined. To study the association of TFIIH before and after bubble rewinding, particle populations with concrete TFIIH features were intersected with particle populations from EC classes and ITC classes, respectively. The identified particles were then subjected to global 3D refinement.

Additionally, particles with features of a rewound bubble were also classified with a mask around the RNA exit tunnel. This resulted in two main species. First, particles with the TFIIB- ribbon occupying the RNA exit tunnel and second, particles with the full-length 14 nt RNA occupying the RNA exit tunnel. Intersection of particles with full-length 14 nt RNA with particles from EC14a resulted in 16,312 mutual particles (25% of EC14a). Intersection of particles with full-length 14 nt RNA with particles from EC14b resulted in 25,046 mutual particles (31% of EC14b). This suggests that the competition between RNA and TFIIB-ribbon in the RNA exit tunnel is independent of TFIIE as particles with full-length 14 nt RNA were similarly distributed in the two classes of different TFIIE states. Particles with the full-length 14 nt RNA were then subjected to focused refinement with a mask around core Pol II. The final composite maps were created with the focused refined map of core Pol II and EC14a/b.

### Model building

To facilitate model building, cryo-EM maps were filtered according to local resolution in Relion3.1^68,69^ and auto-sharpened in Phenix^70,71^. For modelling of the ITCs, previous published models (PDB: 7nw0^38^, 5flm^61^) were rigid-body docked into focused refinement maps in Chimera^72^ and adjusted manually in Coot^73^. The resulting models were then real-space refined in Phenix^70,71^ followed by consecutive rounds of rebuilding with Coot^73^ and ISOLDE^74^. Final models showed good stereochemistry as validated by Molprobity^75^. For modelling of EC14a/b, core Pol II (without stalk) from previous published model (PDB: 5flm^61^) was rigid-body docked into focused refinement maps of core Pol II in Chimera^72^ and adjusted manually in Coot^73^. The resulting models were then real-space refined in Phenix^70,71^ followed by consecutive rounds of rebuilding with Coot^73^ and ISOLDE^74^. Afterwards, TFIIE, Pol II stalk from previous published model (PDB: 7nw0^38^) and core Pol II were rigid-body docked into focused refinement maps in Chimera^72^ and real-space refined in Phenix^70,71^. The real-space refined models and TFIIF from a previously published model (PDB: 7nw0^38^) were rigid-body docked into focused refinement maps in Chimera^72^ to generate the final models of ECs. The final EC models were subjected to comprehensive validation (cryo-EM) in Phenix^70,71^ and showed good stereochemistry.

## Quantification and statistical analysis

Signals of RNA transcripts from abortive transcription assays were quantified with ImageJ 1.53^76^. Abortive transcription experiments were performed four times independently. The percentage of aborted RNA was defined as: (transcription signals from fractions 1-4)/(transcription signals from fractions 1-4+7.5*transcription signals from fractions 7-22). Statistical significance of the results was calculated with Brown-Forsythe ANOVA tests using GraphPad Prism 9.5.1. Ultracentrifugation binding experiments were performed twice.

## SUPPLEMENTARY FIGURE LEDENDS

**Figure S1.**
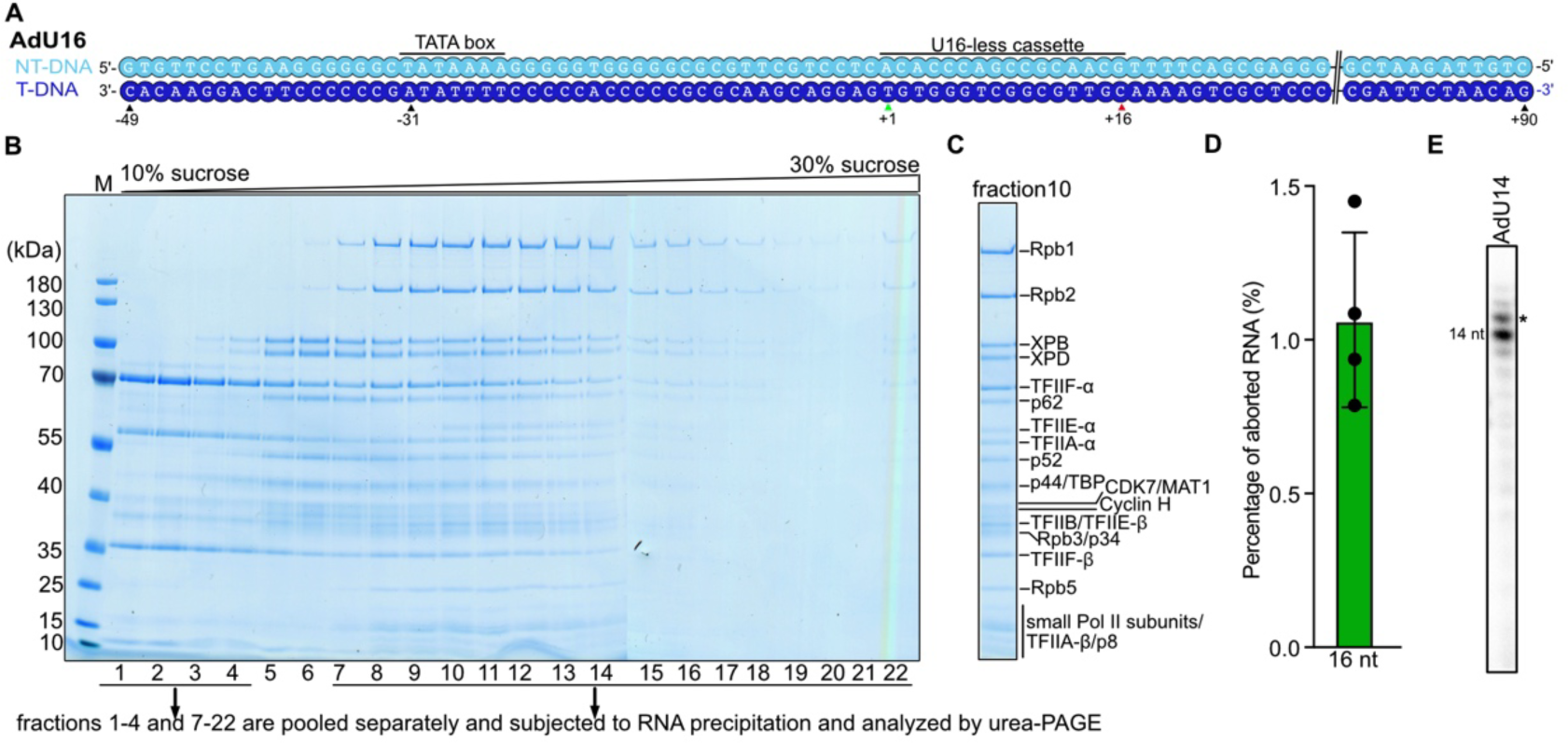
Transcription reactions with AdU16 and AdU14 templates. **A,** Scheme of AdU16 template. The TSS (+1) is highlighted by a green triangle. Pol II paused site (+16) is highlighted as red triangle. The register of DNA is indicated with respect to the TSS (+1). **B,** SDS-PAGE analysis of fractions 1-22 after sucrose gradient ultracentrifugation. Protein complexes generated from transcription reactions with AdU16 and AdU14 templates migrate similarly on a 10-30% sucrose gradient (data not shown). An exemplary gel from a transcription reaction with the AdU14 template is shown. Fractions containing Pol II (7-22) and without Pol II (1-4) are pooled separately and subjected to isopropanol precipitation. M: marker. **C,** Protein identification of fraction 10. **D,** Quantification of the percentage of aborted transcript from the transcription reaction with the AdU16 template. The experiments were repeated four times. **E,** Urea-PAGE analysis of *in vitro* transcription assays performed with the AdU14 template. Pol II tends to mis-incorporate an additional nucleotide when it reaches the end of U-less cassette. The transcript resulted from misincorporation is labeled with an asterisk. The assay was repeated twice.

**Figure S2.**
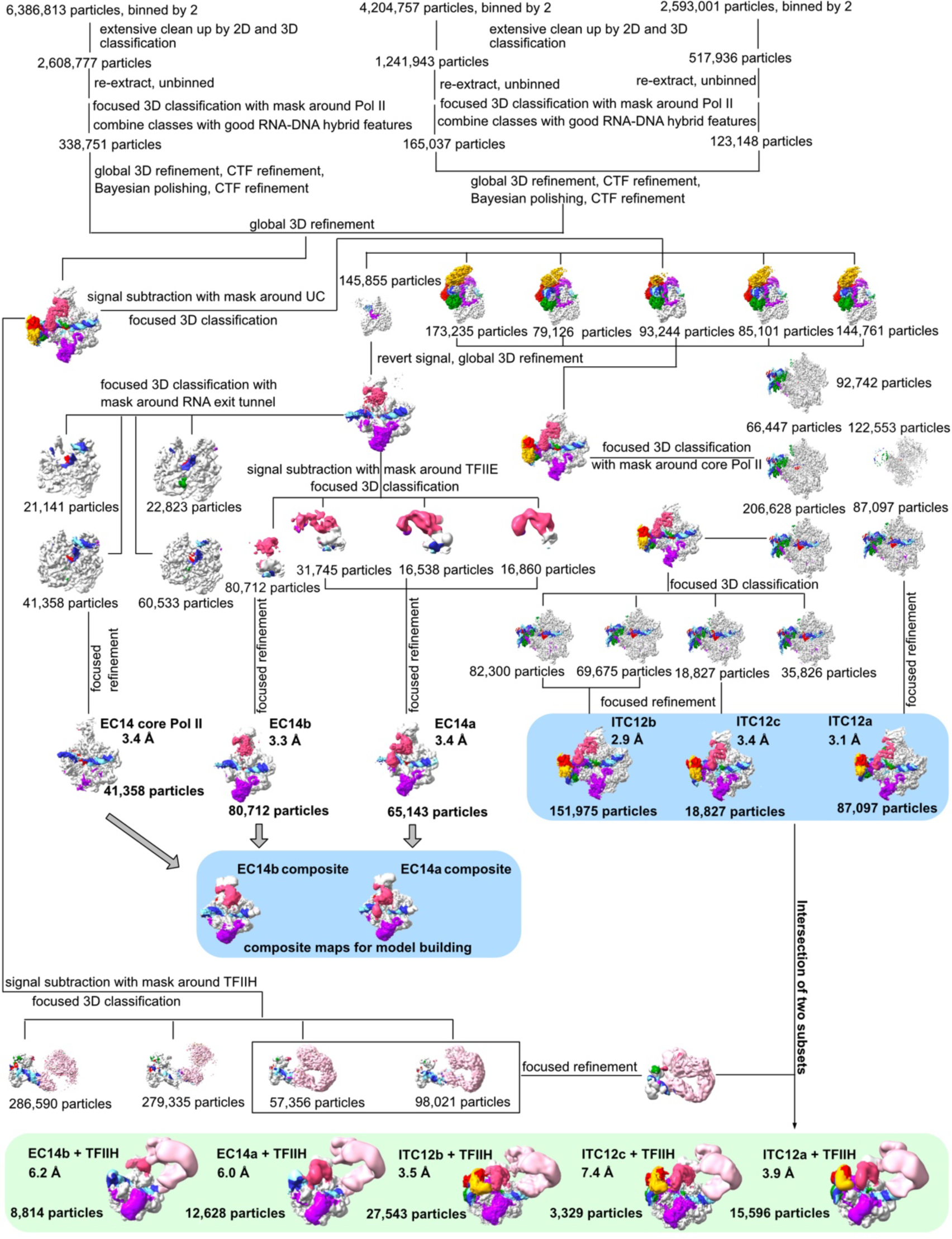
Cryo-EM analysis of ITC12a-c and EC14a/b. Cryo-EM data processing scheme. Reconstructions used for model building are depicted in blue boxes. Reconstructions from intersected particles with both features for TFIIH and for Pol II are depicted in the green box. The final particle number and resolution of each reconstruction are reported next to the densities.

**Figure S3.**
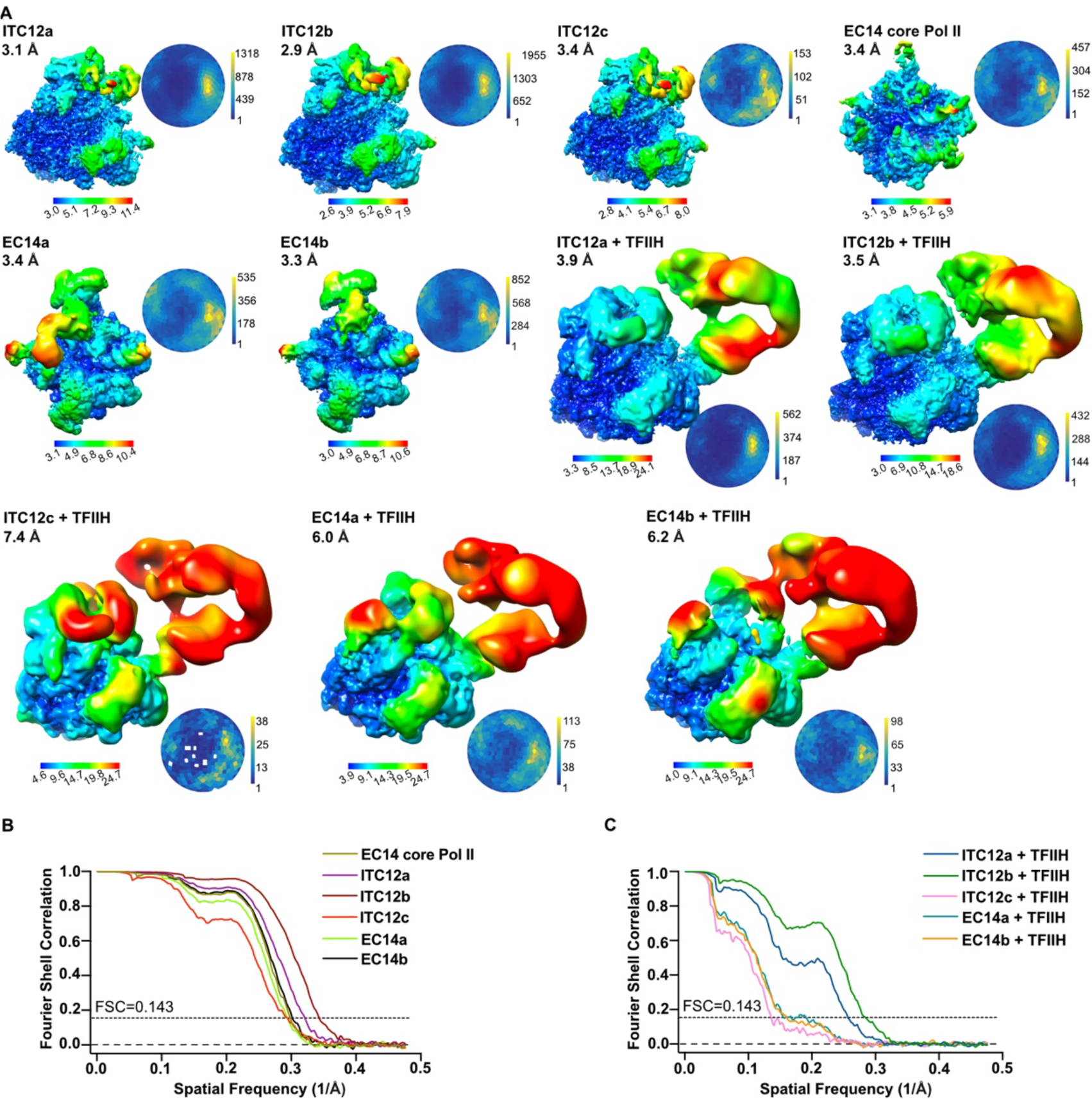
Quality of the cryo-EM reconstructions of transcription complexes obtained with AdU14 template. **A,** All focused refined maps and global refined maps are filtered according to local resolution. Global resolution and angular distribution of each map are shown next to the densities. **B**, Fourier shell correlation (FSC) for focused refined maps. Global resolutions reported in **A,** are estimated at the FSC 0.143 cut-off criterion. **C**, Fourier shell correlation for globally refined maps of reconstructions containing TFIIH. Global resolutions reported in **A,** are estimated at the FSC 0.143 cut-off criterion.

**Figure S4.**
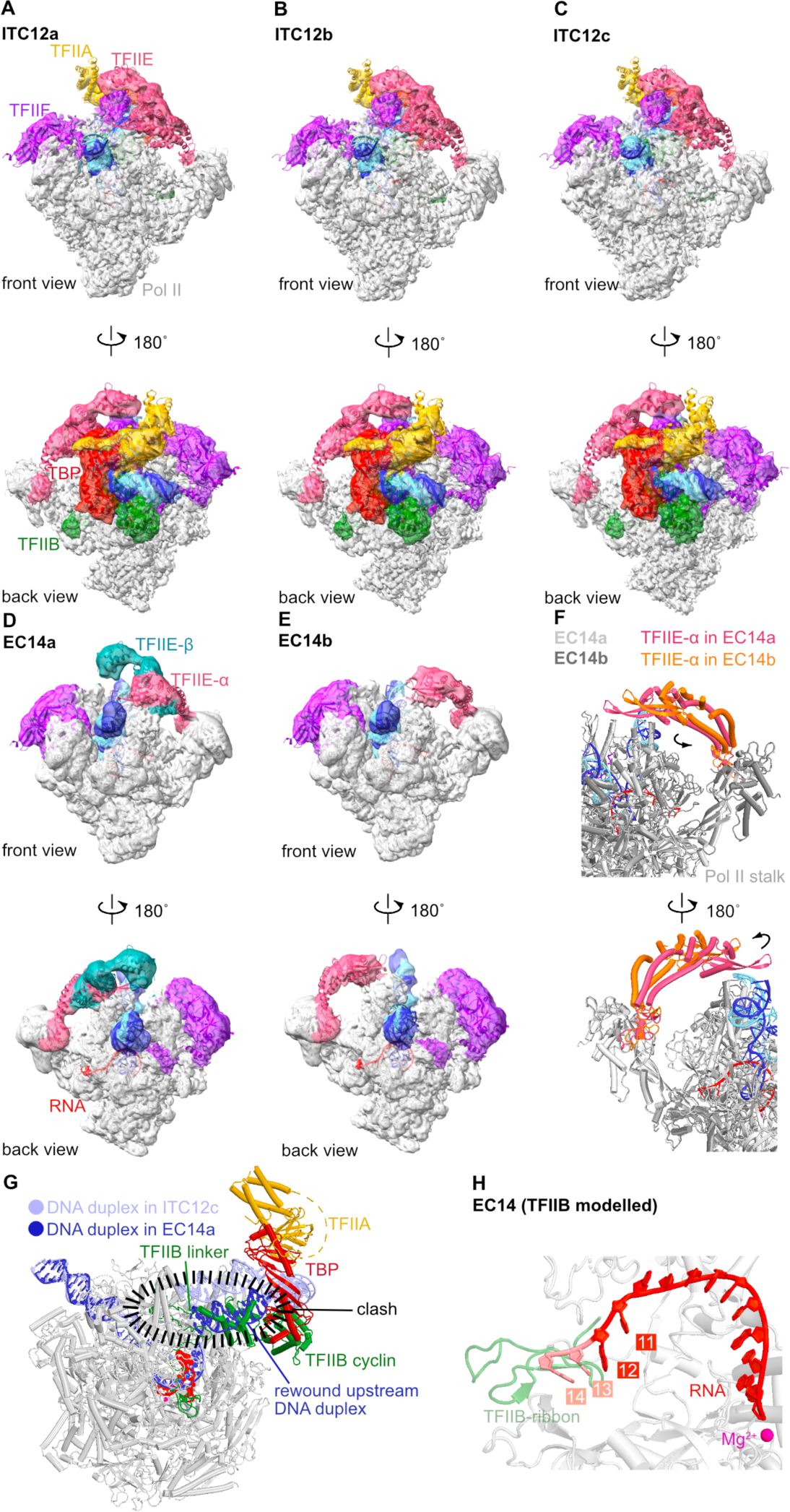
Cryo-EM maps and structures of ITC12a-c and EC14a/b. **A-E,** Cryo-EM maps and structures (docked into maps) of ITC12a-c and EC14a/b. Two views are shown for each structure. **F,** Zoom-in on TFIIE and Pol II of EC14a and b. Pol II of EC14a is colored in light grey. Pol II of EC14b is colored in dark grey. TFIIE-α of EC14a is colored in pink. TFIIE-α of EC14b is colored in orange. The structures of EC14a and EC14b are superimposed using Rpb1. **G,** Superposition of ITC12c and EC14a reveals clashes between the rewound upstream DNA duplex with TFIIB linker and cyclin domains. For clarity, TFIIE, TFIIF and TFIIH are omitted. **H,** Eviction of TFIIB ribbon by RNA. The TFIIB ribbon clashes with the RNA at length of 13-14 nt and is displaced in EC14. The first two RNA bases that would clash with the TFIIB ribbon in EC14 are colored in salmon.

**Figure S5.**
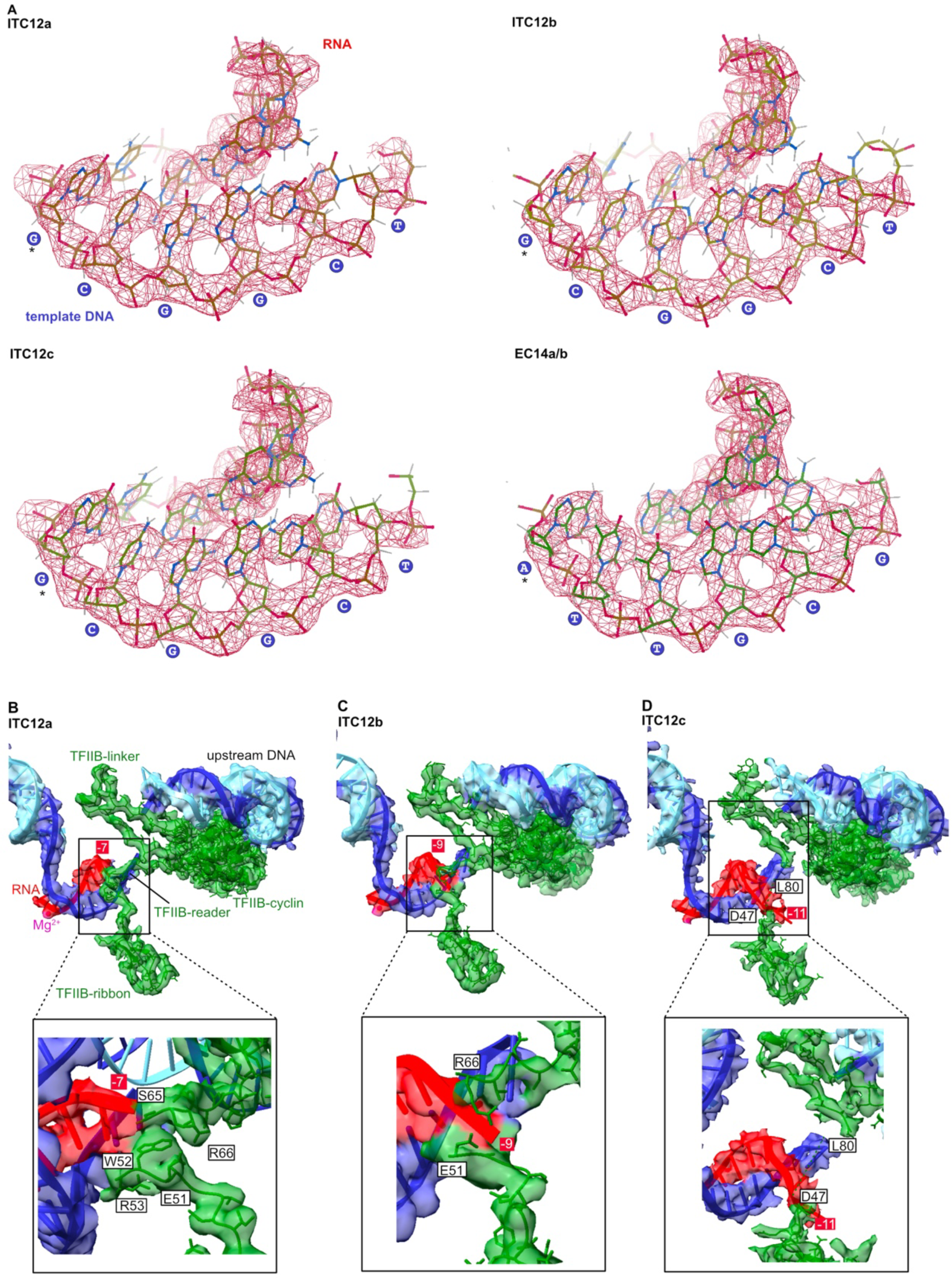
Close-up views of the densities for the DNA-RNA hybrids of ITC12a-c and EC14a/b and details of nucleic acids-TFIIB interactions in ITC12a-c. **A,** Cryo-EM densities of the DNA-RNA hybrids are shown. The map resolutions allow distinguishing purines and pyrimidines in the template DNA strand. The sequence of the template DNA is shown and the template base for register -1 is highlighted with an asterisk. **B-D,** Cryo-EM maps and structures of TFIIB and the nucleic acids in ITC12a-c are shown. The position of the first visible base at 5’-end of the RNA relative to the active site (-1) is marked. The active site Mg^2+^ is shown as a magenta sphere.

**Figure S6.**
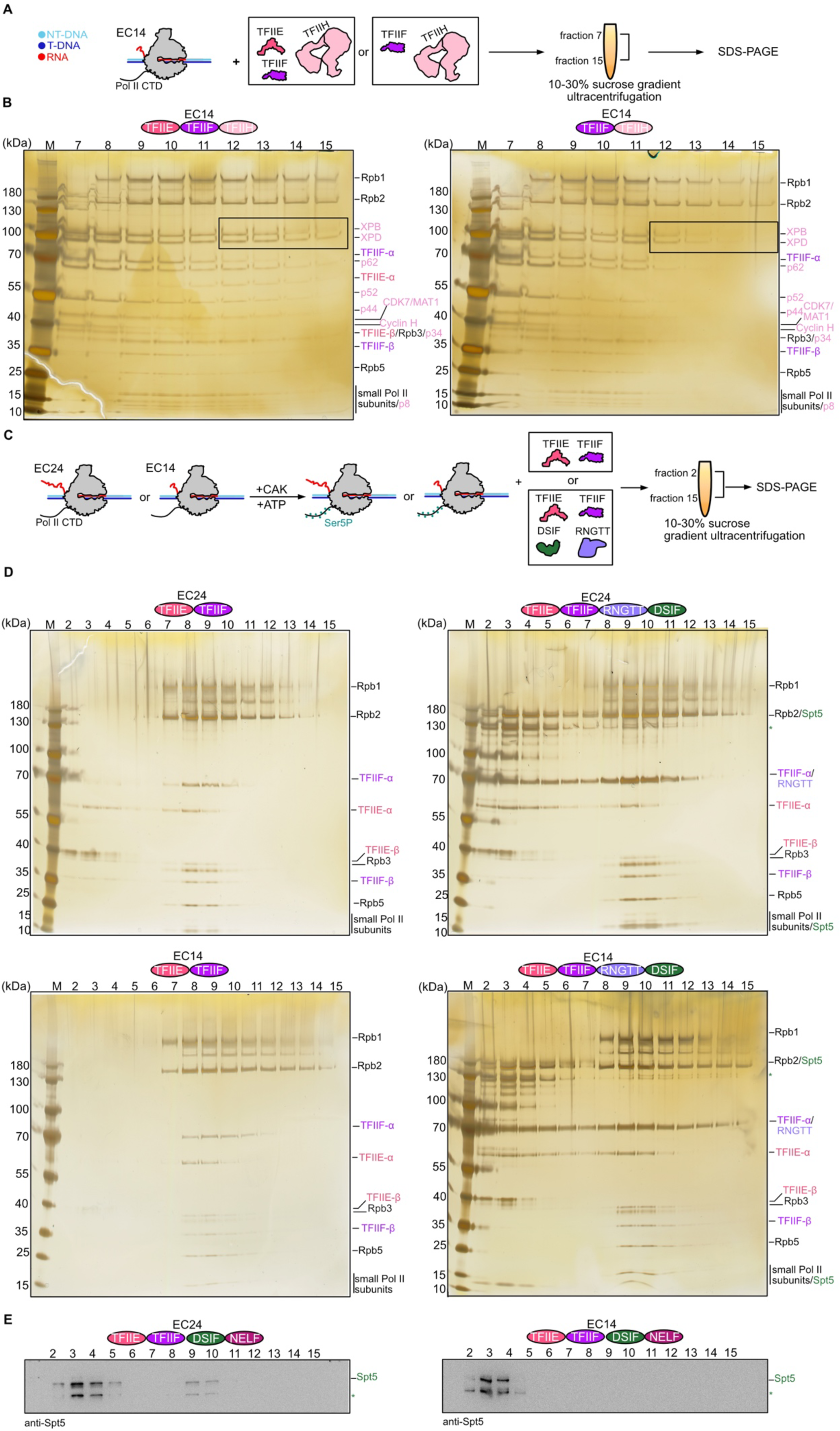
Biochemical analysis of the displacement of TFIIH and TFIIE from the early EC. **A,** Scheme of the experimental setup to monitor the displacement of TFIIH. For details, see Methods. **B,** SDS-PAGE analysis of peak fractions after analytical gradient ultracentrifugation. TFIIH migrated into later fractions and formed a complex with Pol II only in the presence of TFIIE. The experiment was repeated twice. **C,** Scheme of experimental setup to monitor the displacement of TFIIE by DSIF and RNGTT. For details, see Methods. **D,** SDS-PAGE analysis of peak fractions after analytical gradient ultracentrifugation with the experimental setup as in **C**. The experiment was repeated twice. Green asterisk marks a degradation band of Spt5. **E,** Western blot analysis of Spt5 from peak fractions after analytical gradient ultracentrifugation with the experimental setup as in **Figure 5C**. Green asterisk marks the degradation band of Spt5. The experiment was repeated twice.

**Figure S7.**
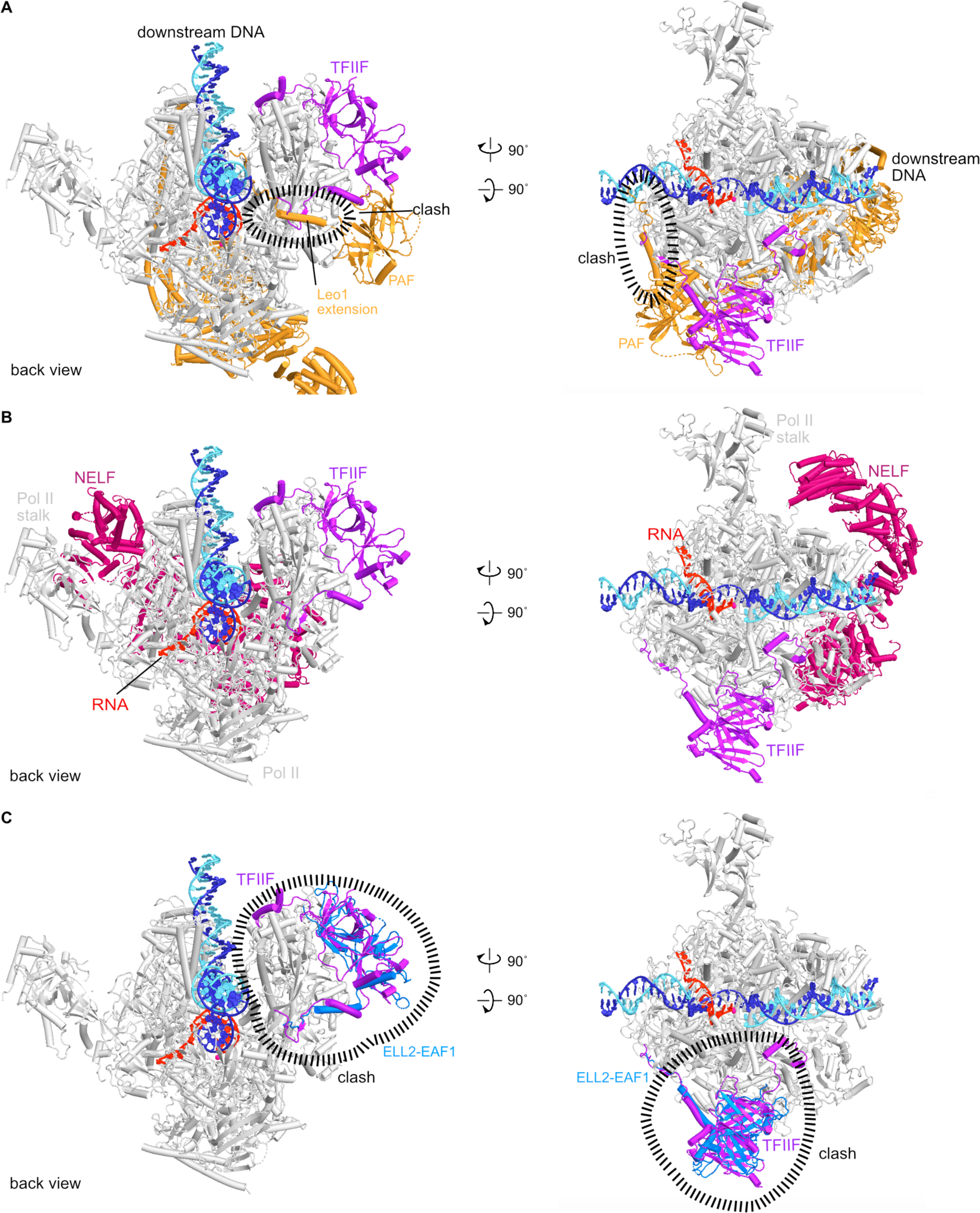
Structures of EC14-TFIIF and comparison with NELF, PAF and ELL2-EAF1 **A,** Superposition of the EC14-TFIIF structure onto the activated EC (PDB: 6gmh^25^). For clarity, DSIF and SPT6 are omitted. TFIIF clashes with the Leo1 extension of PAF. **B,** Superposition of the EC14-TFIIF structure onto the paused EC (PDB: 6gml^35^). For clarity, DSIF is omitted. TFIIF does not clash with NELF. **C,** Superposition of the EC14-TFIIF structure onto Pol II-DSIF-ELL2-EAF1 (PDB: 7oky^26^). For clarity, DSIF is omitted. TFIIF clashes with ELL2-EAF1, which is part of the SEC.

## SUPPLEMENTARY DATA

**Supplementary Video 1 | Three-step model for promoter escape by mammalian Pol II.**

## KEY RESOURCES TABLE

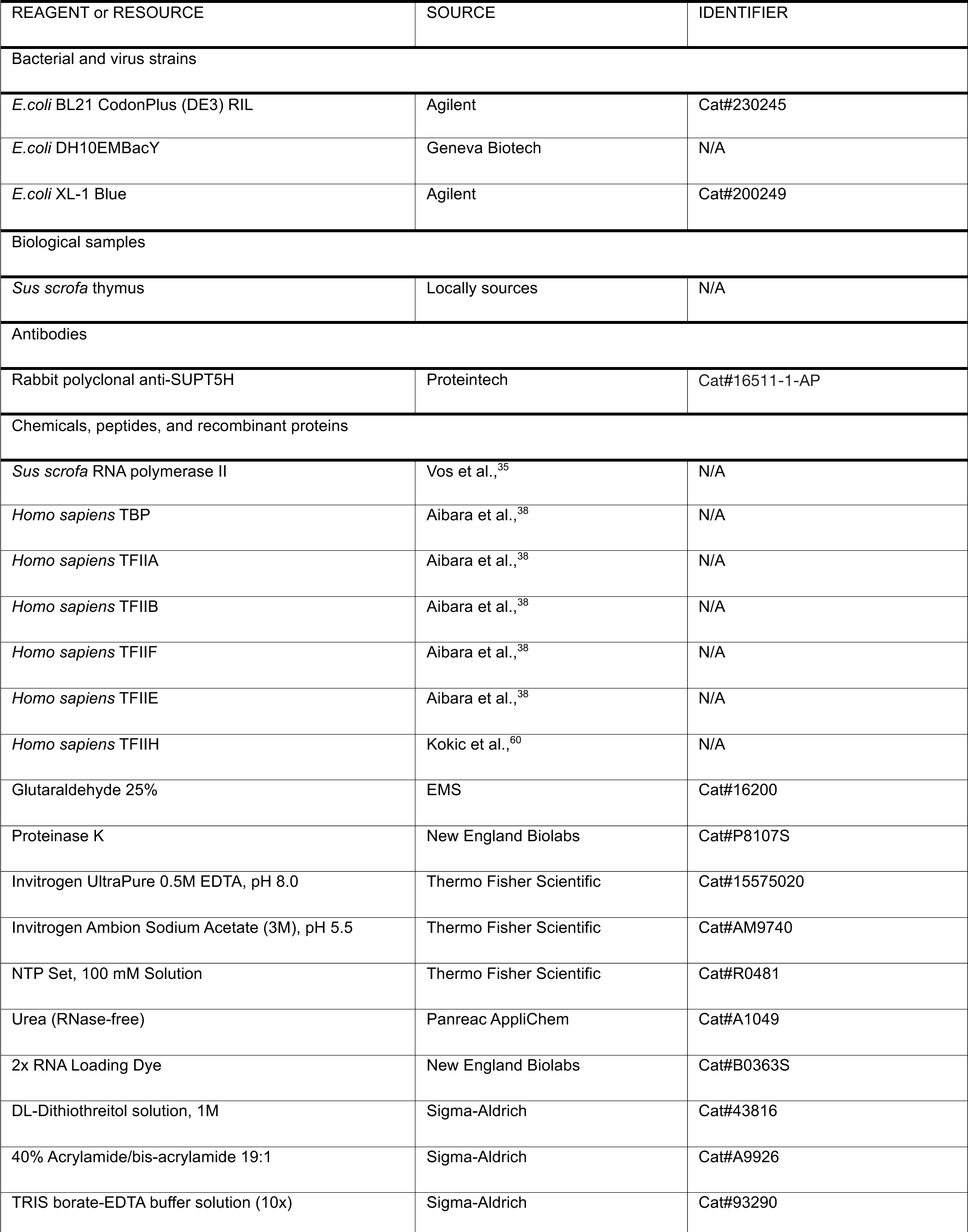
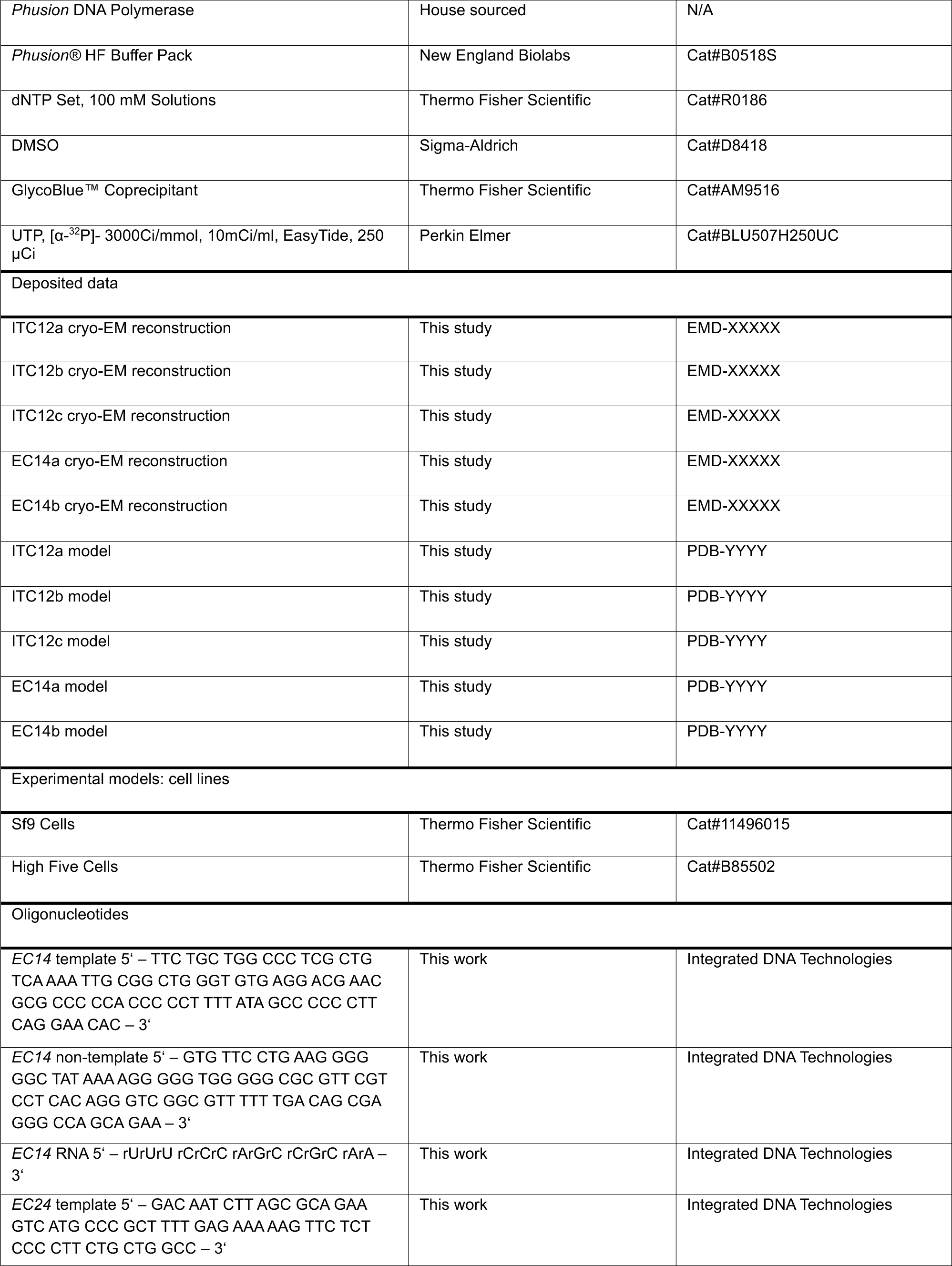
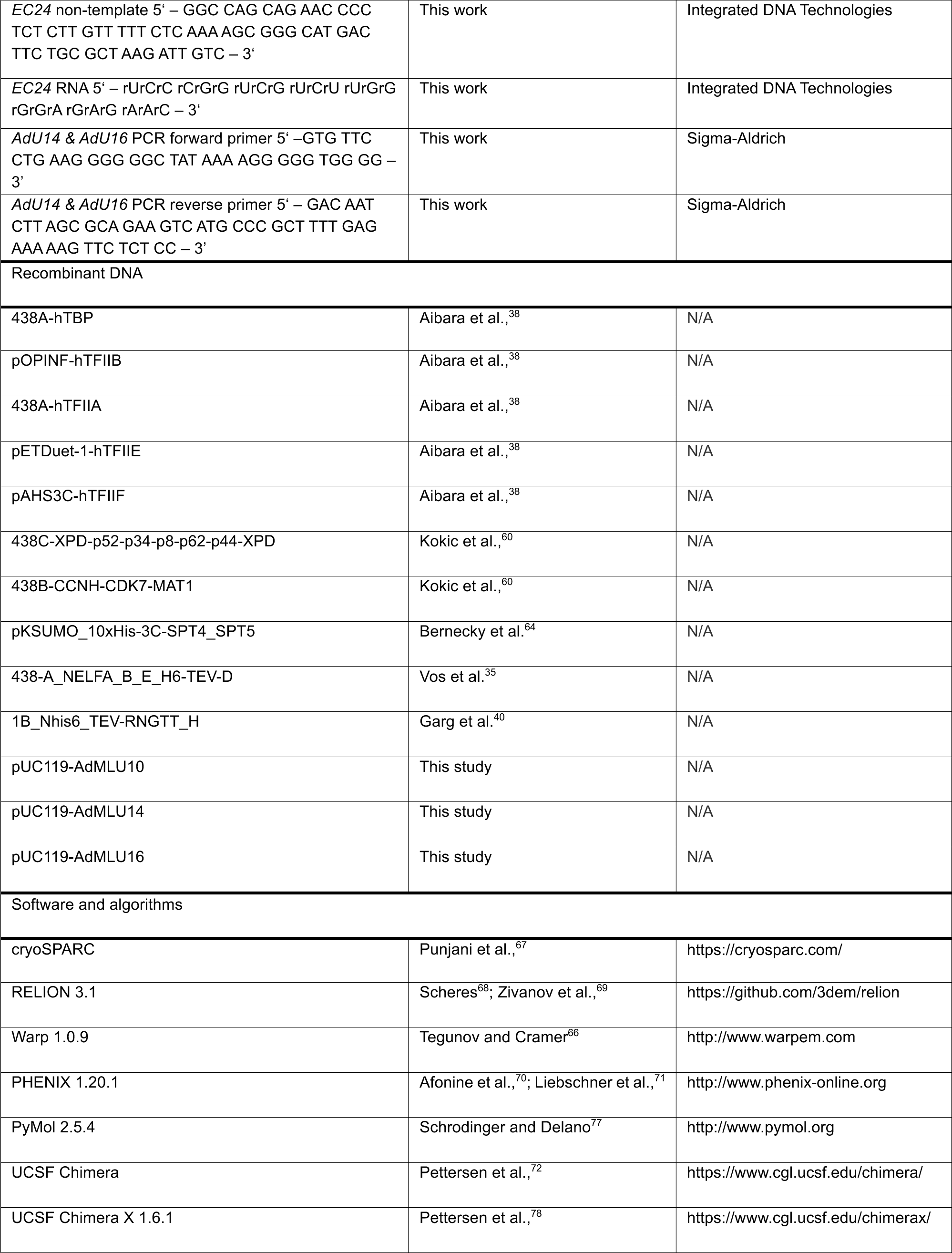
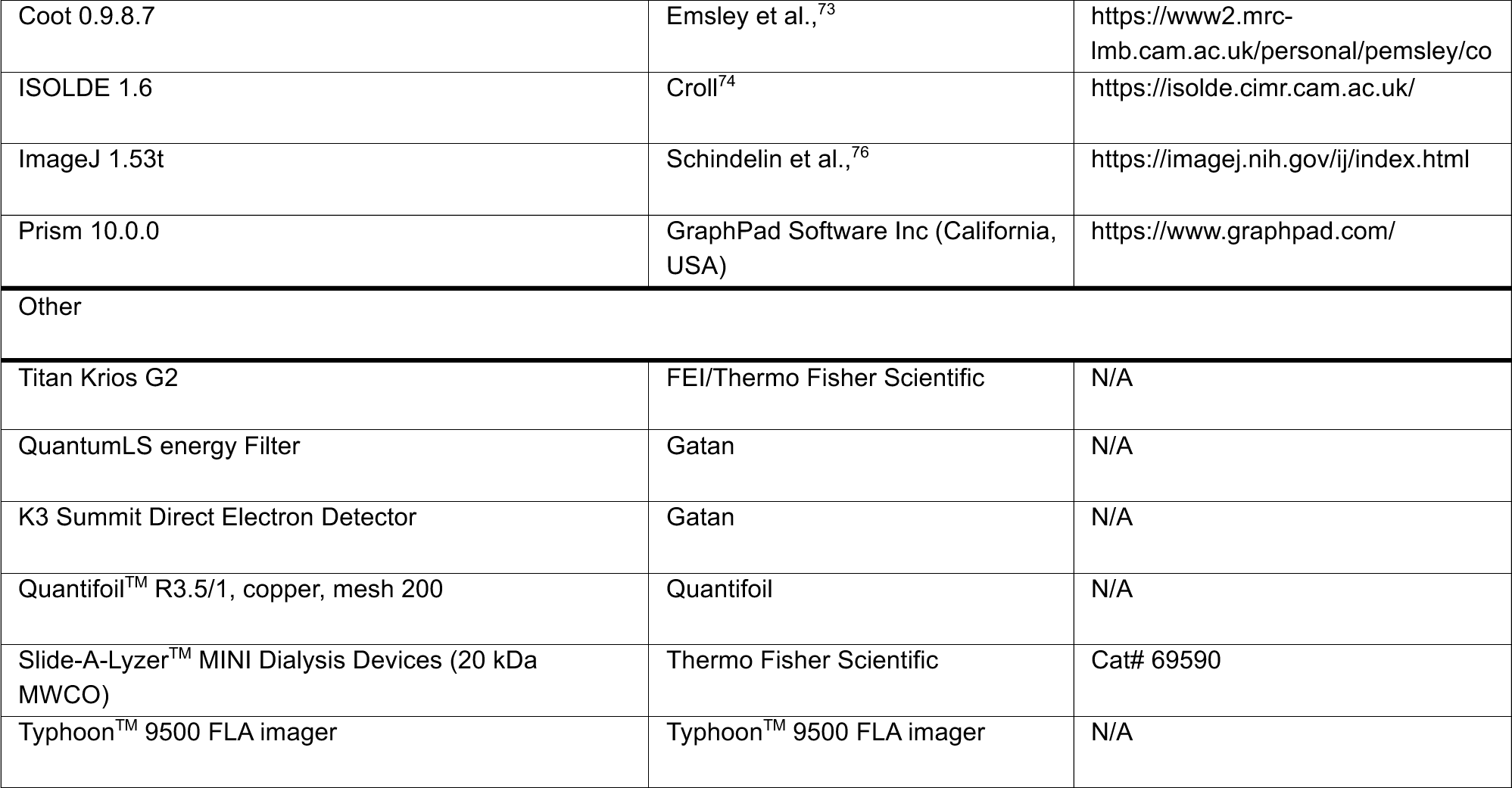

## Notes

### Competing Interest Statement

The authors have declared no competing interest.

